# Frame-shift mutation of *InCO* might cause early flowering of Japanese morning glory and might have contributed to northward expansion

**DOI:** 10.1101/2024.09.05.611556

**Authors:** Hiroaki Katsuyama, Takuro Ito, Kyousuke Ezura, Emdadul Haque, Atsushi Hoshino, Eiji Nitasaka, Michiyuki Ono, Shusei Sato, Sachiko Isobe, Hiroyuki Fukuoka, Nobuyoshi Watanabe, Tsutomu Kuboyama

## Abstract

Japanese morning glory (*Ipomoea nil*), a short day plant, has been used for studying flowering times. Here, quantitative trait loci (QTL) analysis for days from sowing to flowering (DTF) of F_2_ between *I. nil* var. Tokyo Kokei Standard (TKS) and *I. hederacea* line var. Q65, an early flowering variety, revealed four QTLs: *QTL Ipomoea Flowering 1–4* (*qIF1–4*). The position of *qIF3*, which had the most significant effect among the four QTLs, corresponds with that of *I. nil* (or *I. hederacea*) *CONSTANS* (*InCO*/*IhCO*) in the linkage map. There is a single-base In/Del in the coding sequence of *InCO*/*IhCO*. The single-base deletion (SBD) causes a frame-shift mutation and loss of function in TKS allele (*inco-1*). *I. nil* accessions bearing *inco-1* tend to flower early, similarly to rice varieties bearing the loss of function allele of *CO* ortholog, *hd1*. Consequently, *inco-1* was inferred to reduce DTF. This inferred effect of *inco-1* corresponds with the effect of the TKS allele of *qIF3*. Because *inco-1* is found exclusively in Asian accessions, the SBD in *inco-1* might have played an important role in the expansion of Japanese morning glories, originally native to the tropical regions of the Americas, into temperate Asia.

## Introduction

Japanese morning glory (*Ipomoea nil*), a short-day (SD) plant that is highly sensitive to the photoperiod, has been used as a model plant to study flowering (**Hayama *et al*. 2007**, **Higuchi *et al*. 2011, Takeno 2016**). Genetic pathways that promote flowering in response to seasonal cues were first reported in the model species *Arabidopsis thaliana* (**Andres and Coupland 2012**), in which *CO* expression is regulated by light. The circadian clock plays a central role in regulating photoperiodic flowering (**Andres and Coupland 2012**). *CO* encodes a zinc finger transcription factor, which binds to the promoter of *FLOWERING LOCUS T* (*FT*), a florigen gene, and which positively affects *FT* expression (**Cao *et al*. 2014, Putterill *et al*. 1995**).

For SD plants, intensive genetic studies of flowering have been conducted in rice (**Hori *et al*. 2016**). Several genes key to controlling the heading date of rice have been identified using quantitative trait locus (QTL) analysis. They have been cloned. *Heading date 1* (*Hd1*), the ortholog of Arabidopsis *CO*, is a major gene controlling the heading date in rice (**Yano *et al*. 2000**). In fact, *Hd1* regulates expression of the *Heading date 3a* (*Hd3a*) rice florigen gene, delays heading under LD, and promotes heading under SD (**Hayama *et al*. 2003**, **Kojima *et al*. 2002, Tamaki *et al*. 2007, Yano *et al*. 2000**).

From the study of *I. nil* flowering, the homologous genes of *Arabidopsis* flowering genes have been isolated (**Hayama *et al*. 2007, Higuchi *et al*. 2011, Liu *et al*. 2001, Zheng *et al*. 2009**). The *I. nil CO* ortholog, INIL11g18779, was originally named *PnCO* because of the former scientific name of Japanese morning glory: *Pharbitis nil*. As presented herein, the prefix "*Pn*" of genes derived from *I. nil* is replaced with "*In*". Therefore, *PnCO* will be referred to as *Ipomoea nil CONSTANS* (*InCO*) herein. *InCO*, which is highly expressed in SD condition, has multiple splicing variants: a transcript containing a long intron (*InCO* (li)), a transcript containing a short intron (*InCO* (si)), and a transcript containing no intron (*InCO* (ni)) (**Liu *et al*. 2001**). Only the *InCO* (ni) mRNA encodes complete CONATANS-like protein with a CO, CO-like TOC1 (CCT) domain. It can restore an *Arabidopsis co* mutant. By contrast, *InCO* (li) and *InCO* (si) encodes truncated protein lacking the terminal CCT domain (**Liu *et al*. 2001**). Two *I. nil* homologous genes of *FT*, *PnFT1* and *PnFT2*, expressed only in inductive SD conditions and overexpression of *PnFT1*, promote flowering in *Arabidopsis* and *I. nil* (**Hayama *et al*. 2007)**. Transgenic plants constitutively expressing *I. nil* ortholog of *GIGANTEA*, *PnGI*, form fewer flowers than non-transgenic plants do. However, the expression level of *InCO* does not differ dramatically in transgenic plants compared to non-transgenic plants (**Higuchi *et al*. 2011**). Consequently, although the homologous genes of flowering related genes in *Arabidopsis* have been reported in *I. nil*, their function in photoperiodic flowering remains unclear.

The whole genome sequence of *I. nil* ‘Tokyo Kokei Standard’ (TKS) has been sequenced. Useful genome sequences are now available (**Hoshino *et al*. 2016a**). EST-SSR marker of *I. nil* has also been reported: approximately half of them are polymorphic between TKS and Ivy-leaved morning glory, *I. hederacea* ‘Q65’ (Q65) (**Ly *et al*. 2012**). In addition, single nucleotide polymorphisms (SNP) become easily detected by comparing mRNA sequences between two accessions using the next generation sequencing (NGS) (**Edward *et al*. 2011**). Consequently, these techniques enable us to map QTLs for days from sowing to flowering (DTF) in morning glory.

Actually, *I. nil* has varieties of various flowering times (**Yoneda and Takenaka 1981**). Consequently, natural variations in *I. nil* are attractive resources for mapping QTLs affecting flowering time in dicot SD plants. Flowering in late July in Ibaraki, Japan, TKS is a standard variety of Japanese morning glory. Q65 was probably transported from the United States to Japan along with grains. It blooms approximately two weeks earlier than TKS. This report describes QTL mapping of DTF in the F_2_ population between Q65 and TKS and a frame-shift mutation found in the TKS allele of *InCO*, which is the candidate gene of the most prominent QTL of DTF.

## Materials and Methods

### Plant materials

*I. nil* accessions TKS (Q1065), Q31, Q33, Q61, Q62, Q63, and Q1187 and an *I. hederacea* accession, Q65, and the F_2_ population derived from the interspecific cross Q65 × TKS were provided by the Morning glory stock center of Kyushu University with support by the National Bio-Resource Project of the MEXT, Japan. An *I. nil* accession, PI227365 was provided by Agricultural Research Service, United States Department of Agriculture. *I. nil* varieties, Pekin-tendan (PKT) and Yakuyou-shirohana (YYS) were derived from the genetic stock of Ibaraki University (**Hoshino *et al*. 2016b**). An *I. nil* variety, Violet, was purchased from Marutane Seed Co., Kyoto, Japan. After dividing 192 individuals from the F_2_ population, each of the 96 individuals was cultivated in 2011 and 2012. These plants were cultivated under natural conditions at the College of Agriculture, Ibaraki University (36.36°N, 140.12°E), Ami, Ibaraki, Japan.

### SSR marker development from I. nil and I. batatas

Using 10% acrylamide gel electrophoresis 326 SSR markers from *I. nil* (**Ly *et al*. 2012**) and 1250 EST-SSR markers from *I. batatas* (Kazusa DNA Lab, Japan) were analyzed. Polymorphic markers between TKS and Q65 were selected for genotyping the F_2_ population (**Supplemental Table 1**). DNA was extracted from fresh leaves of each plant using a DNeasy Plant Mini kit (Qiagen Inc.). Each 6 µl of reaction mixture contained 1xNH_4_ reaction buffer (Bioline; Meridian), 0.2 mM dNTP, 3 mM MgCl_2_, 0.2 µM each primer, 0.15 unit *Taq* polymerase (Bioline; Meridian) and 1.0 ng of template DNA. Polymerase chain reactions (PCR) were run using a modified ‘touchdown PCR’ program (**Sato *et al*. 2005**).

### Development and genotyping of SNP markers

After mRNA was isolated from 0.1 g buds of Q65 using an RNeasy Plant mini kit (Qiagen Inc.), it was sequenced using a Genetic Analyzer II (Illumina Inc.). Using software (Maq; **Li *et al*. 2008**), the sequence reads were mapped to a reference sequence, the non-redundant *I. nil* ESTs derived from TKS buds cDNA (**Hoshino *et al*. 2016a, Ly *et al*. 2012**), and were used to detect SNPs between Q65 and TKS. The RNA-seq data were archived at the DNA Data Bank of Japan under accession number DRA010196. These SNPs were genotyped by high-resolution melting (HRM) analysis (**Liew *et al*. 2004**) or melting temperature (T_m_)-shift primer method (**Fukuoka *et al*. 2008, Wang *et al*. 2005**). Primers for HRM analysis were designed with Primer 3 (**Rozen and Skaletsky 2000**) (**Supplemental Table 2**). In designing of primers for HRM, Primer 3 parameters of PCR-amplicon length, optimal melting temperature, and primer length were set respectively as 80–110 bp, 60°C, and 20 bases. PCR amplification and DNA melt curve analysis were performed in a total volume of 10 µl containing 10 mM Tris-HCl (pH 8.3), 65 mM KCl, 1.5 mM MgCl_2_, 0.2 µM each primer, 0.2 mM dNTP, 1.25% glycerol, 0.4× Eva Green (Biotium Inc.), 0.25 U *Taq* DNA polymerase (Ampliqon AS) and 1.0 ng of template DNA (**Fukuoka *et al*. 2008**).

Primers for T_m_-shift primer method were designed using ’tms_primer_designer.pl’ (**Fukuoka *et al*. 2008**) (**Supplemental Table 3**). The PCR for Tm-shift primer method was performed in Quantification / DNA binding Dye / Standard curve mode of an Eco real-time PCR system (Illumina Inc.). The PCR conditions were 94°C for 3 min, 40 cycles of 95°C for 20 s, 58°C for 30 s, and 72°C for 30 s with subsequent melt curve analysis using the default settings of the system. The PCR for HRM analysis was performed in High Resolution Melt/DNA binding Dye/DNA/PCR with HRM Curve mode of the Eco real-time PCR system (Illumina Inc.). The PCR conditions were 94°C for 3 min, 40 cycles of 96°C for 20 s, 58°C for 1 min, and 72°C for 30 s followed by melt curve analysis using the default settings of the system.

### Linkage map construction and QTL analysis

From genotyping data of the F_2_ population, we constructed linkage maps using QTL IciMapping software ver. 4.2 (**Meng *et al*. 2015**). DTF was calculated from the date of sowing to the date on which the first flower of an individual plant opened. Using each constructed linkage map and DTF-trait data, we detected QTLs for DTF with R/qtl package (**Broman *et al*. 2003**). We conducted interval mapping, composite interval mapping, and multiple QTL analysis, respectively using the function, "scanone", "cim" and "stepwiseqtl". To establish alpha = 0.05 significant thresholds, the data were permutated 1000 times in each experiment. The estimated percent variances explained for the QTL and effects of QTL were calculated using the function "fitqtl".

### PCR amplification and DNA sequencing of the Q65 ortholog of InCO (IhCO)

Four primer sets were used for PCR amplification of *IhCO* and its 5′ flanking region (**Supplemental Table 4**). PCR amplifications were performed using KOD FX Neo DNA polymerase (Toyobo Co. Ltd., Osaka, JAPAN). The nucleotide sequences of the PCR products were analyzed with a capillary sequencer (ABI3130; Life Technologies Inc.) using the primers presented in **Supplemental Table 4**. The DNA sequence data derived from *IhCO* were archived at the DNA Data Bank of Japan under accession nos. LC716650 and LC768960. The alignments of nucleotide sequences and putative amino-acid sequences were found using Clustal W ver. 2.1 (**Larkin *et al*. 2007**). Splicing site prediction was performed with NetGene2-2.42 server (**Hebsgaard *et al*. 1996**, https://services.healthtech.dtu.dk/services/NetGene2-2.42/).

### Quantitative reverse transcription PCR (qRT-PCR) analysis

TKS and Q65 plants were cultivated at 25°C under continuous light (195 µmol m^-2^s^-1^) until five days after sowing and were grown under SD (10 h light/14 h dark) or LD (14 h light /10 h dark) conditions for 3 days. Then, cotyledons were harvested after treating 8 h or 14 h of dark period from the last light period. Total RNA was isolated from cotyledons using an SV Total RNA Isolation System (Promega Corp.). cDNA was synthesized using Prime Script RT Master mix (Takara Bio Inc.). Real-time PCR was conducted using a Light Cycler 96 system (Roche) with TB Green^®^ *Premix Ex Taq*™ II (Takara Bio Inc.). Three primer sets, ni (forward primer (Fwd) 5′- AGGGACTGCAGCAGCATAAC-3′ and reverse primer (Rev) 5′- TGGAGACCATATCCCGTGTT-3′), si (Fwd 5′- CCATCAGTCACACAGTCTCCA-3′ and Rev 5′- TGAAGCTGTGGAGGCATCTG-3′) and li (Fwd 5′- TCCAATAAAACCCAACGTCA-3′ and Rev 5′-GGGGAAGTTGCATACCTTGA-3′), were designed and used for qRT-PCR analyses of three *InCO*/*IhCO* splicing variants. *InACTIN4* (Fwd 5′-GAATACTTGTATGCCACGAGCA-3′ and Rev 5′-GGATTGCCAAGGCAGAGTAT-3′) was used as the internal reference gene. Real-time PCR conditions were 94°C for 3 min, 40 cycles of 95°C for 10 s, 58°C for 30 s, and 72°C for 15 s, followed by melt curve analysis using default settings of the system. Relative expression levels for each sample were calculated based on the comparative Ct method. Welch’s two-tailed *t*-test was used to compare the expression levels between TKS and Q65.

## Results

### DTF in F_2_ populations

In 2011 and 2012, seeds of the F_2_ population were sown respectively on May 20 and June 7. The DTF values of individuals were measured (**Fig. 1**): those for Q65 and TKS were, respectively, 54 and 62 days in 2011 and 38 and 52 days in 2012. The ranges of DTF in the F_2_ populations of 2011 and 2012 were 46–80 days and 39–76 days, respectively; more than half of individuals flower later than both parents (**Fig. 1**).

**Fig. 1.**
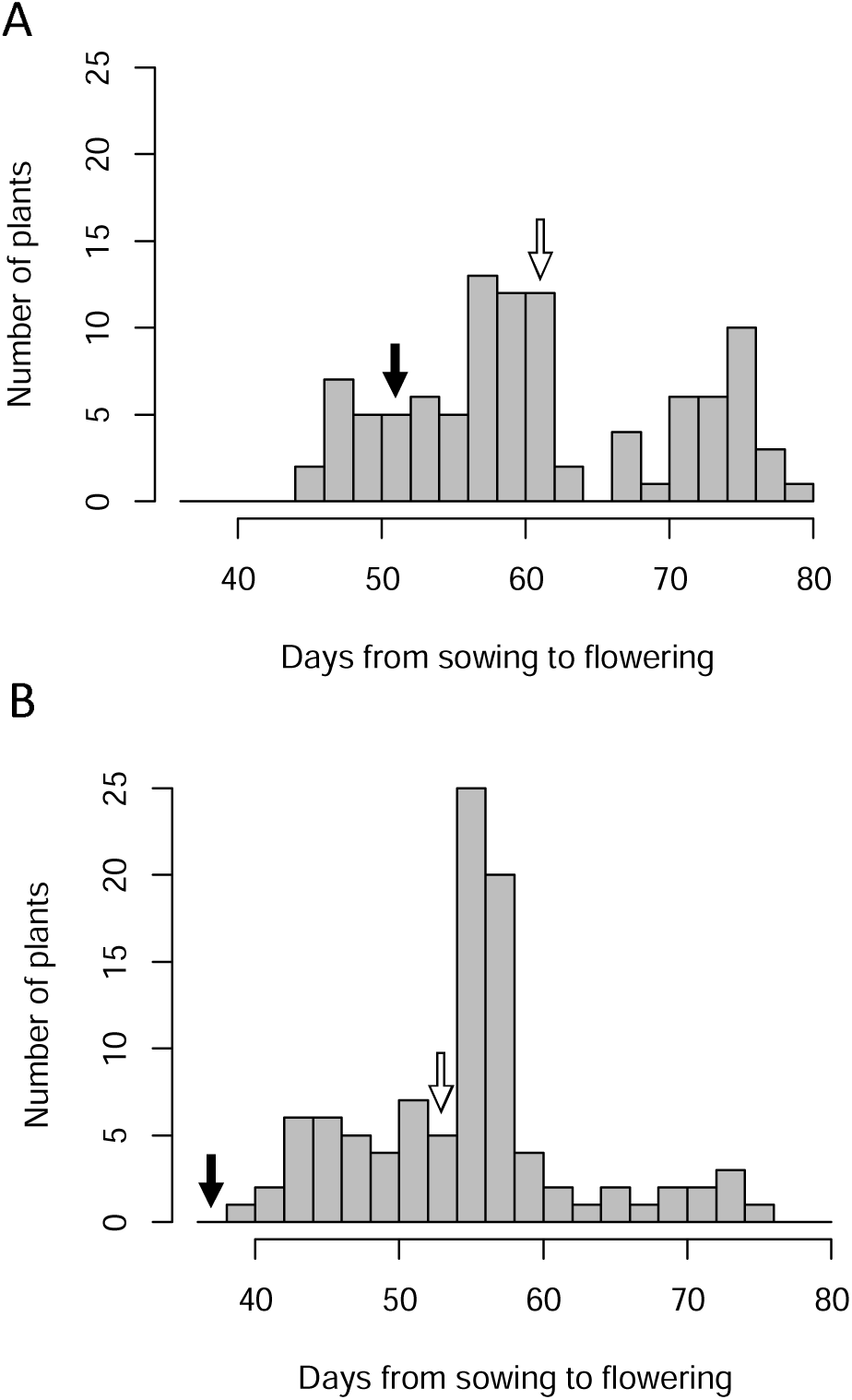
Distribution of days from sowing to flowering among F_2_ population derived from a cross: Q65 × TKS. A and B respectively show the F_2_ population in 2011 and 2012. The black arrow and white arrow respectively indicate flowering days of Q65 and TKS.

### Marker development and linkage-map construction

EST-SSR markers derived from *I. nil* (**Ly *et al*. 2012**) and *I. batatas* (**Supplemental Table 1**) showing polymorphisms between Q65 and TKS in polyacrylamide gel electrophoresis were used for linkage-maps construction. For SNP marker development, sequence reads of Q65 buds mRNA were mapped to the non-redundant ESTs derived from *I. nil* (**Hoshino *et al*. 2016a, Ly *et al*. 2012**). The SNPs detected between Q65 and TKS were used for developing SNP markers. An F_2_ population of 96 individuals derived from the cross between Q65 and TKS was cultivated in 2011 and 2012. They were genotyped by DNA markers for constructing genetic linkage maps. The linkage map of the 2011 population contained 254 loci including 53 Tm-shift-SNP markers, 39 HRM-SNP markers, 146 EST-SSR markers of *I. nil*, 16 EST-SSR markers of *I. batatas* and two phenotypic markers (**Supplemental Figs. 1–2, Supplemental Tables 1–3**). The linkage map covered a total length of 1902 cM and included 15 linkage groups. The linkage map of the 2012 population included 187 loci and covered a total length of 1736 cM and consisted of 15 linkage groups. These constructed linkage groups were anchored to chromosomes using pseudo-chromosomes of Asagao_1.1 (**Hoshino *et al*. 2016a**) **(Supplemental Figs. 1–2**).

### QTL analysis for DTF

In the F_2_ population cultivated in 2011, four QTLs for DTF, designated as *qIF1*, *qIF2*, *qIF3*, and *qIF4*, were mapped in the vicinity of the DNA marker of IES0160, respectively, on chromosome 5, Contig11987 on chromosome 9, Contig4567.156 on chromosome 11, and Contig05575 on chromosome 14 based on a multiple QTL model (**Table 1**). In this QTL model, the phenotype y is modeled as y = *qIF1* + *qIF2* + *qIF3+ qIF4* + *qIF1*:*qIF4* (A colon between two QTLs denotes interaction between the QTLs.). In the F_2_ population cultivated in 2012, a QTL is detected only at the *qIF3* position (**Table 1**). The LOD value of *qIF3* in 2012 was reduced by approximately one-third compared to that in 2011. In 2012, seeds were sown two weeks later than in 2011. The shorter period of long-day condition in 2012 population might reduce power for detection of QTL related to photoperiodism.

**Table 1.**
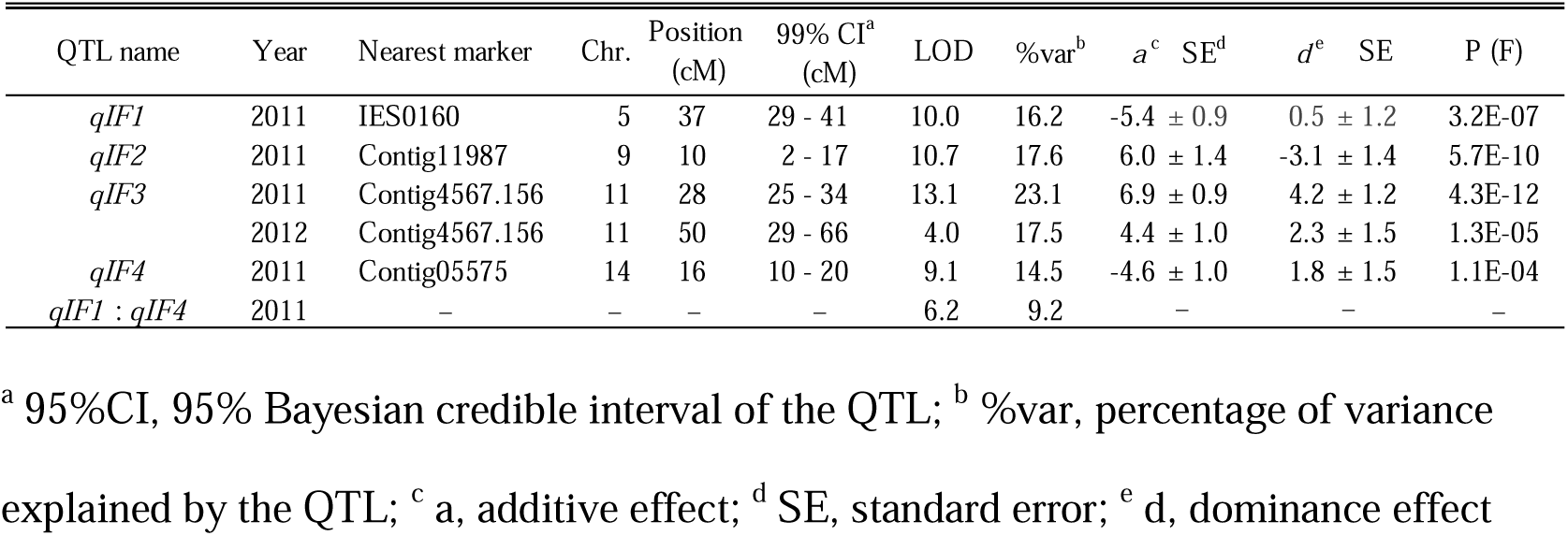
Multiple QTL mapping of DTF in the cross between Q65 and TKS.

Q65 allele of *qIF2* and *qIF*3 increased DTF and delayed flowering (**Fig. 2**). By contrast, the Q65 allele of *qIF1* and *qIF4* decreased DTF and enhanced flowering. The percentage of phenotypic variance explained (PVE) by these QTLs was 14.5% – 23.1%; the total PVE was 73.7% based on the multiple QTL model in R/qtl (**Table 1**). *qIF3* showed the highest PVE among these QTLs (23.1%). These results found for the 2011 population demonstrated the readily apparent effects of *qIF1*, *qIF2*, and *qIF3*, but not of *qIF4* (**Fig. 2**). In fact, *qIF4* was detected neither in the simple interval mapping nor in the composite interval mapping (**Supplemental Fig. 3**). However, *qIF4* had some interaction with *qIF1* (**Supplemental Figs. 4 A, B**). The effects of *qIF4* differed depending on the *qIF1* genotype. In individuals with a homozygous TKS allele of *qIF1*, plants that are homozygous for the TKS allele of *qIF4* exhibited a delayed flowering compared to plants homozygous for the Q65 allele of *qIF4*. However, among individuals with a homozygous Q65 allele of *qIF1*, plants homozygous for the TKS allele of *qIF4* showed shorter DTF than those homozygous for the Q65 allele of *qIF4.* The DTF differences among the genotypes of *qIF4* were less than those in the homozygous TKS allele of *qIF1* (**Supplemental Fig. 4 B**). Another significant interaction (LOD = 7.12 > 6.39 (*P* = 0.05)) was detected between the chromosomal region at 116 cM of chromosome 14 (14@116) and at 123 cM of chromosome 5 (5@123) (**Supplemental Fig. 4 C**). The effects of TKS and Q65 allele of 5@123 reversed depending on 14@116 genotypes. Homozygous TKS allele of 14@116 enhanced flowering in homozygous for TKS allele of 5@123, but delayed flowering in heterozygous and homozygous for Q65 allele of 5@123 (**Supplemental Fig. 4 C**).

**Fig. 2.**
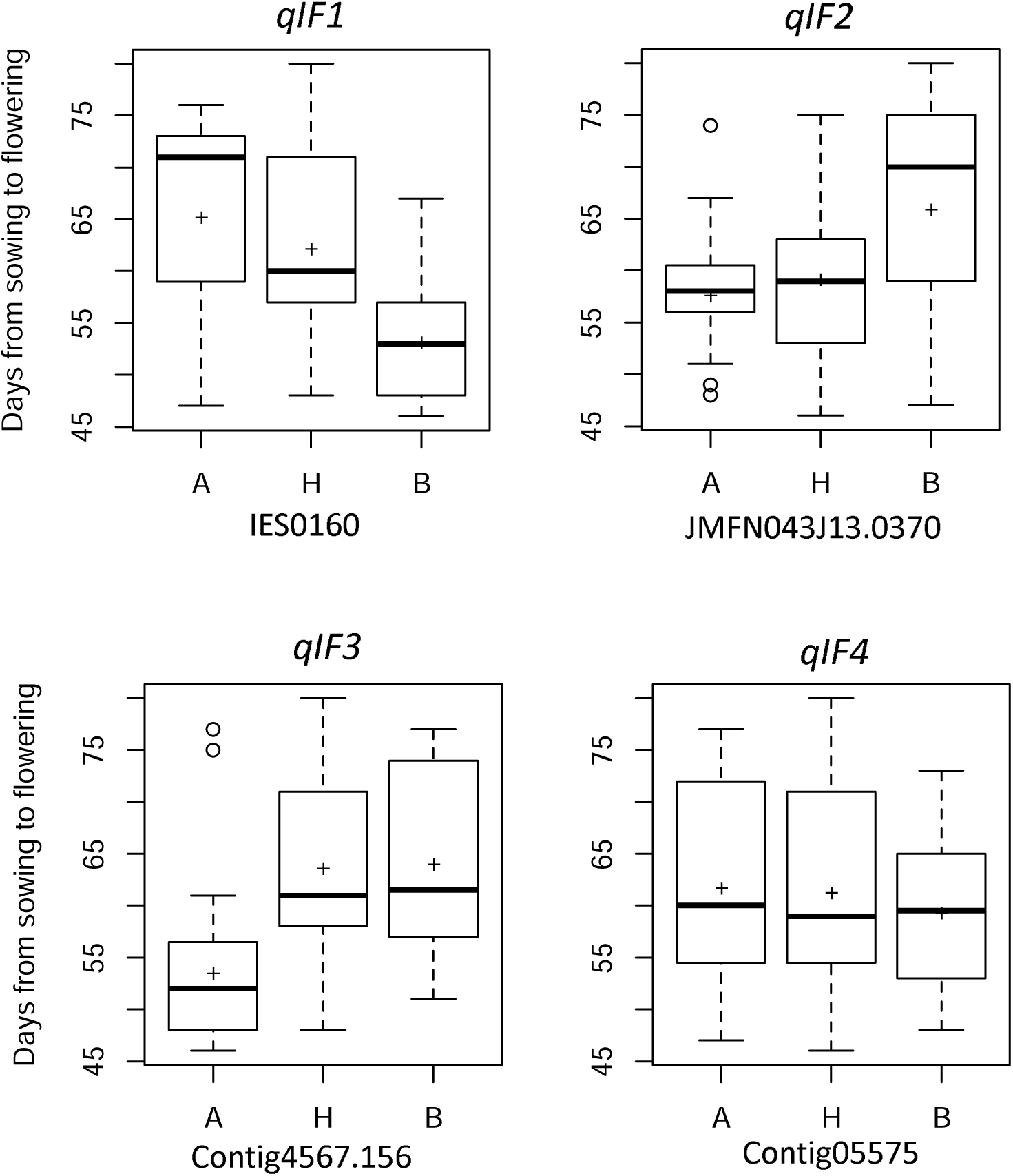
Effects of QTLs genotypes for days from sowing to flowering. A, H, and B of the horizontal axis respectively show TKS allele homozygotes, heterozygotes, and Q65 allele homozygotes. The names of DNA markers are shown under the genotypes.

### Polymorphism between the TKS and Q65 alleles of InCO*/*IhCO

The strongest effect QTLs for DTF, *qIF3* was co-located with *InCO*/*IhCO* in the linkage map. Also, Contig4567.156, the DNA marker the nearest to the LOD peak of *qIF3*, was developed based on polymorphism in the EST encoding *InCO* (**Table 1, Supplemental Figs. 1–2, Supplemental Table 2**). Consequently, *InCO*/*IhCO* might be a candidate of *qIF3*. Therefore, we compared the DNA sequence of *InCO*/*IhCO* between TKS and Q65.

First, using PCR amplification and agarose gel electrophoresis, the 5’ flanking regions of *InCO*/*IhCO* were compared among Q65 and four *I. nil* accessions: PKT, TKS, Q63, and YYS (**Fig. 3**). The size of PCR products amplified from Q65 was 2.3 kbp and approximately 300 bp larger than that of other accessions (**Fig. 3**), which is attributable to the structural difference of the 5′ flanking region of *InCO*/*IhCO* between TKS and Q65. Then, the PCR products of Q65 were sequenced and aligned with the DNA sequence of TKS (**Hoshino *et al*. 2016a**) (**Supplemental Fig. 5**). Short interspersed element (SINE)-like 168 bp insertion sequence was found in the reverse strand of approximately 1.9 kb upstream of the putative transcription start site (**Supplemental Fig. 5**). There were a conserved putative RNA polymerase III promoter containing A-box and B-box (**Galli *et al*. 1981**) and target site duplications (**Fig. 4**). A few thousand copies of this SINE-like sequence were detected by homology search against the Asagao 1.2 genome (**Hoshino *et al*. 2016a**) using the BLAST program. DNA sequences downstream of the SINE-like sequence were highly polymorphic between TKS and Q65. There was a long TA repeat (more than 200 bp) in the Q65 sequence (**Supplemental Fig. 5**).

**Fig. 3.**
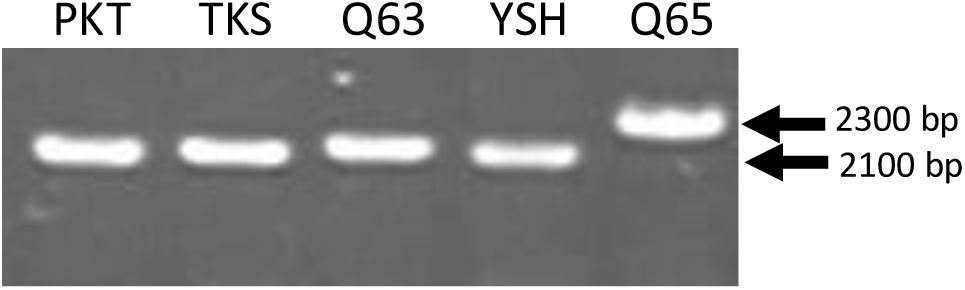
Agarose gel electrophoresis of PCR products amplified from the 5′ flanking regions of *InCO*/*IhCO* in five morning-glory accessions: PKT, Pekin tendan; TKS, Tokyo Kokei standard; YSH, Yakuyo shirohana.

**Fig. 4.**
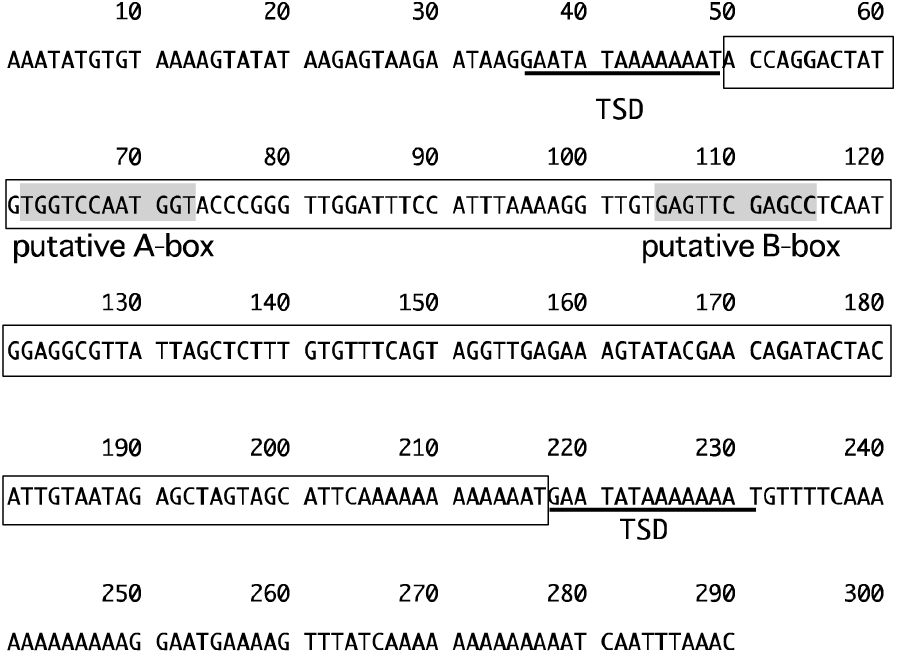
A SINE-like sequence found in the 5′ flanking region of *IhCO.* DNA sequences surrounded by square frame show the SINE-like sequence. TSD, target site duplication. TSDs are shown by underlining. The locations of putative boxA and boxB are shown by a gray background.

In addition to the polymorphisms in the 5′ flanking region, the coding sequence (CDS) of Q65 had 11 single-base substitutions (SBS) and 3 insertions against TKS sequence. Of the 11 SBSs, 8 are synonymous substitutions, which represent functional constraints of *InCO*/*IhCO*. Of the three insertions, the two insertions were 9 and 12 nucleotides, causing an additional 3 and 4 amino acids, respectively, but one was a single-base insertion causing a frame-shift of the translational reading frame (**Fig. 5**). The Q65 allele has a single cytosine base present between positions 7617445 and 7617446 on Chr11 of TKS in the genome sequence (Asagao_1.2) (**Fig. 5**). If the splice sites are identical to *InCO* (ni), the transcript reported for the ’Violet’ allele (**Liu *et al*. 2001**), then the Q65 mRNA would encode an incomplete protein without a CCT domain. Therefore, the splice-donor site of Q65 allele was assessed using NetGene2-2.42 server (**Hebsgaard *et al*. 1996)** (**Fig. 5B**). As a result, the splice-donor site of *InCO* (ni) was not detected confidently as a splice-donor site and 26 nucleotides (nt) downstream from the splice-donor site of *InCO* (ni) was the most confident splice-donor site (**Fig. 5B**. The transcript spliced at 26 nt downstream was correspondent with the transcript reported by Liu *et al*. (**2001**) as *InCO* (si). *InCO* (si) encodes an incomplete protein, which results from premature stop codon in ’Violet’ allele (**Liu *et al*. 2001**), but in the Q65 allele, it encodes complete 433 aa protein with the CCT domain (**Supplemental Fig. 6**).

**Fig. 5.**
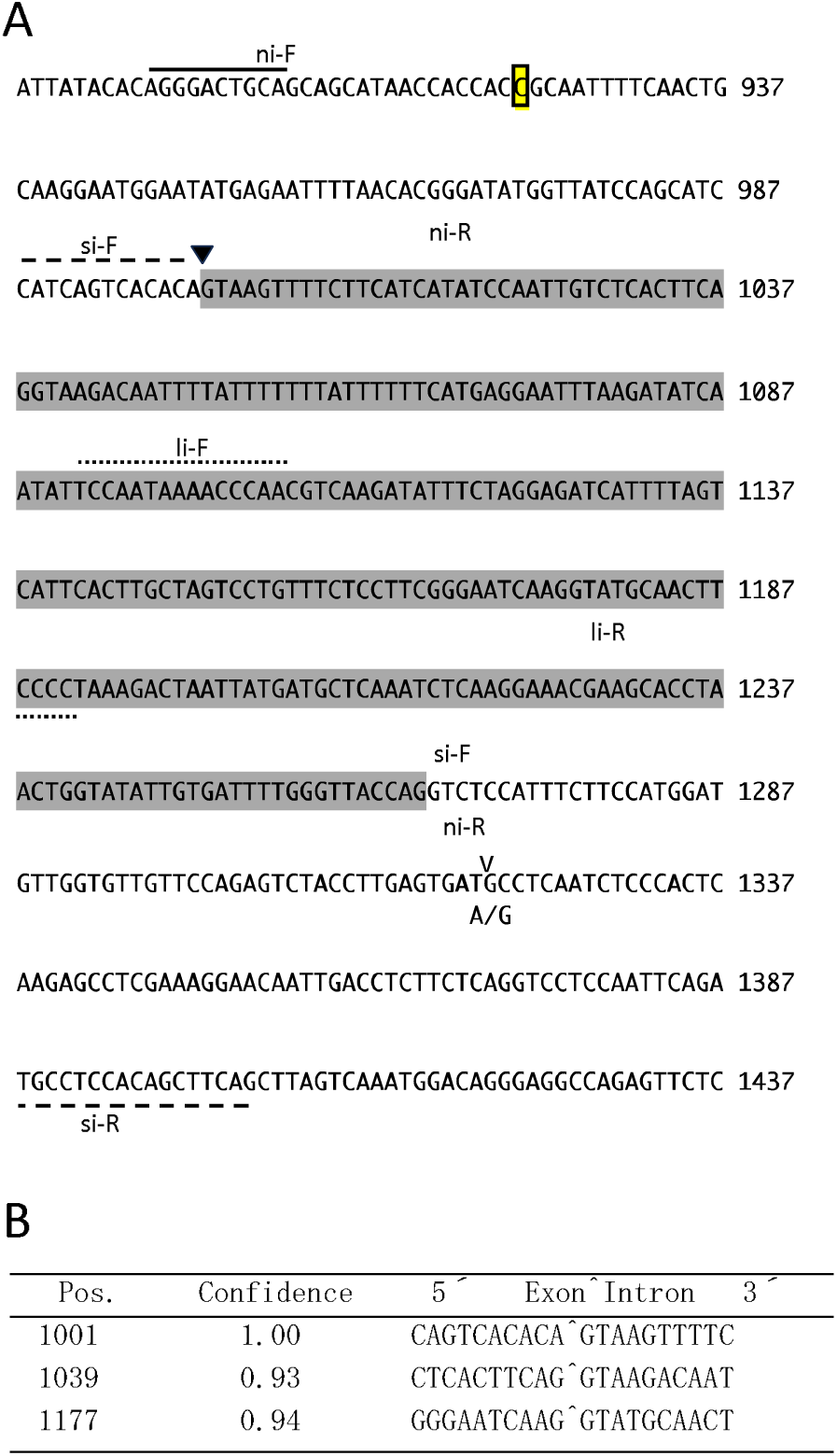
*InCO*/*IhCO*-splice-site prediction at Q65 allele. (A) The genomic DNA sequence around an intron of Q65 allele. Numbers at the right side of DNA sequences show nucleotide numbers from the putative transcription start site. The boxed cytosine is deleted from the TKS allele. A white-reverse triangle shows a splice donor site reported by Liu *et al*. (2001) as no-intron mRNA (*InCO* (ni)). A black reverse triangle shows a splice donor site reported by Liu *et al*. (2001) as short-intron mRNA (*InCO* (si)) and also as predicted by NetGene2 v. 2.4 as the most confident splice-donor site. The DNA sequence with gray background denotes the intron of *InCO* (si). "v" above the DNA sequence signifies the SNP site position. ’A/G’ shows the adenine of TKS allele and the guanine of Q65 allele at the SNP site. Primer pairs used for expression analysis are shown as arrows. (B) A splice-donor-site prediction by Net Gene2 v. 2.4. Pos. denotes the nucleotide position from the putative transcription start site.

### Distribution of the single-base InDel in the CDS of InCO among accessions derived from natural populations

To determine the ancestral form of the single-base InDel in the CDS of *InCO*, the single-base InDel sites were investigated in eight *I. nil* accessions derived from natural populations in addition to TKS and Q65 (**Table 2**. Accessions Q33 and Q1187, derived from the South American continent, have cytosine insertion at the single-base InDel site as Q65 (**Table 2**). Accessions that have the same sequence as TKS in the single-base InDel are limited to those from the Asian countries (**Table 2**). *I. nil* originated from the tropical regions of the Americas (**Austin *et al*. 2001**). The Q65-type sequence of the single-base InDel is apparently ancestral. Moreover, the fact that *InCO* orthologs of *Ipomoea* species *I. triloba* and *I. trifida* also contain the Q65-type sequence in the single-base InDel also support the idea that the Q65-type sequence of the single-base InDel is ancestral (**Fig**. **6**). These findings suggest that TKS type allele (*inco-1*) emerged from the ancestral allele of *InCO* (*InCO-2*) by the single-base deletion (SBD) in the CDS (**Fig. 7A**).

**Fig. 6.**
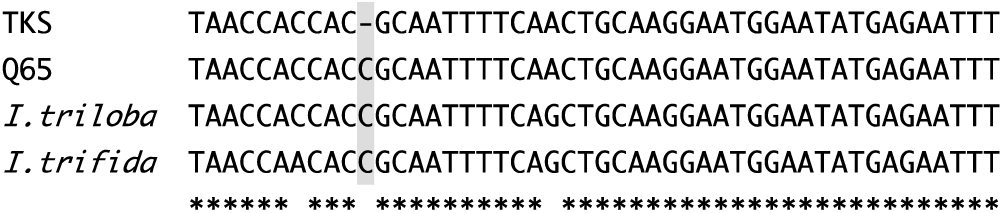
DNA sequence alignment of *InCO* orthologous genes around the single-base InDel causing frameshift in *InCO*. DNA sequence of *I. triloba* (XR_004098766.1) and of *I. trifida* (CP025648.1) were aligned with Clustal W (Ver. 1.83, 2003). Gray background denotes the single-base-InDel site.

**Fig. 7.**
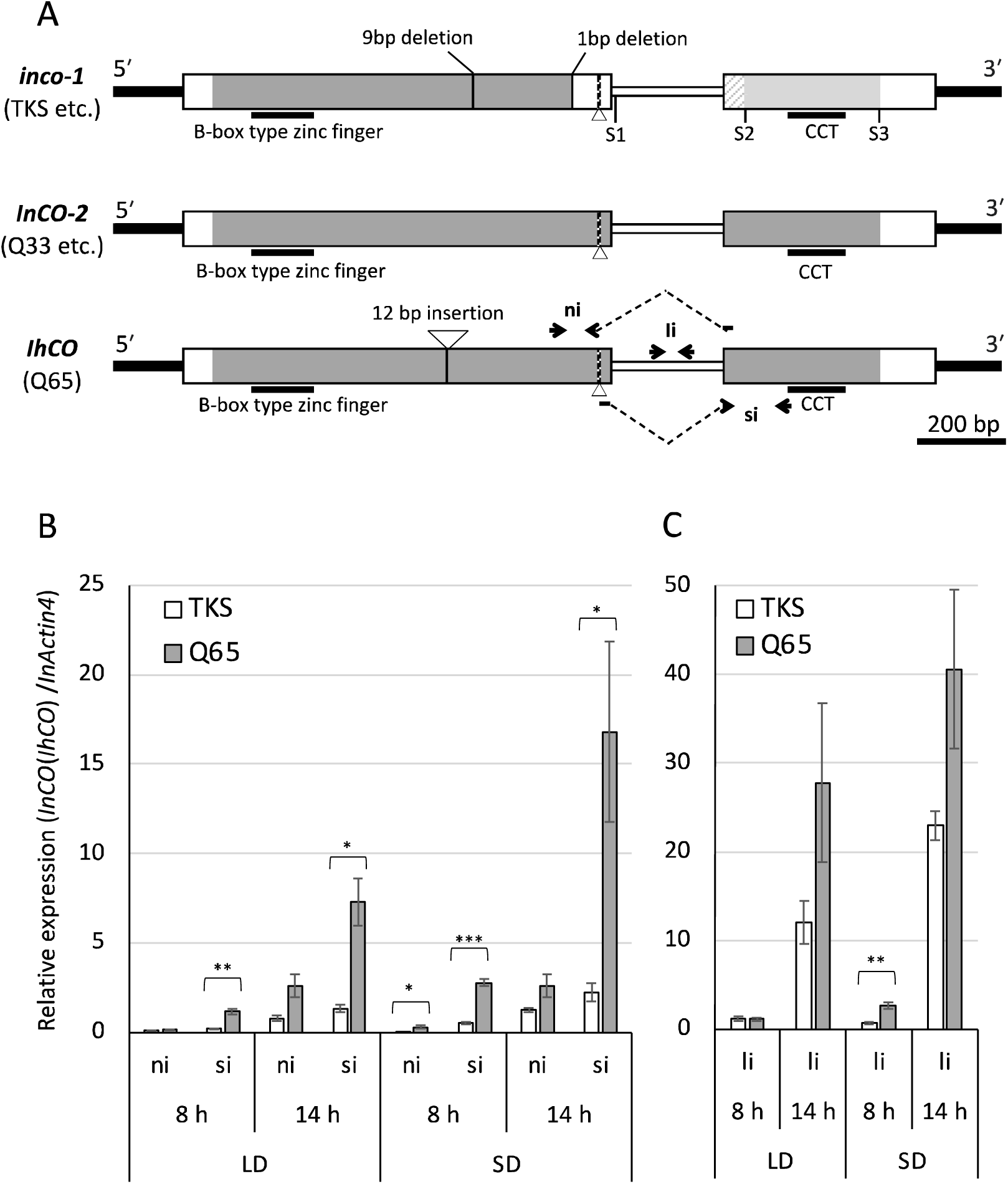
*InCO*/*IhCO* alleles and comparison of the expression levels of three transcripts of *InCO*/*IhCO* in Q65 and TKS. (**A**) Map of the *InCO*/*IhCO* alleles and the PCR primer pairs. Narrow black boxes show untranscribed regions. Wider boxes show exons. A narrow white box shows an intron. White triangles and vertical broken lines show cryptic splice sites producing *InCO* (ni). S1, S2, and S3 of *inco-1* respectively show stop codon sites of *InCO* (li), *InCO* (si), and *InCO* (ni). Gray parts of the boxes show regions homologous to IhCO protein in a.a. Light-gray parts of *inco-1* show that InCO (ni) protein is homologous to IhCO protein. The diagonal stripe pattern shows the region where the InCO (ni) protein is homologous with the IhCO protein, but not with the InCO (si) protein. **ni**, **si**, and **li** indicated by arrows and broken lines are primer pairs used for qRT-PCR. These **ni**, **si**, and **li** were used respectively for amplifying splicing variants, *InCO* (ni), *InCO* (si), and *InCO* (li). (**B, C**) Relative expression analysis of three *InCO* splicing variants using qRT-PCR. *, **, and *** respectively denote significant difference at *P*<0.05, *P*<0.01, and *P*<0.001 by Welch’s two-tailed *t*-test).

**Table 2.**
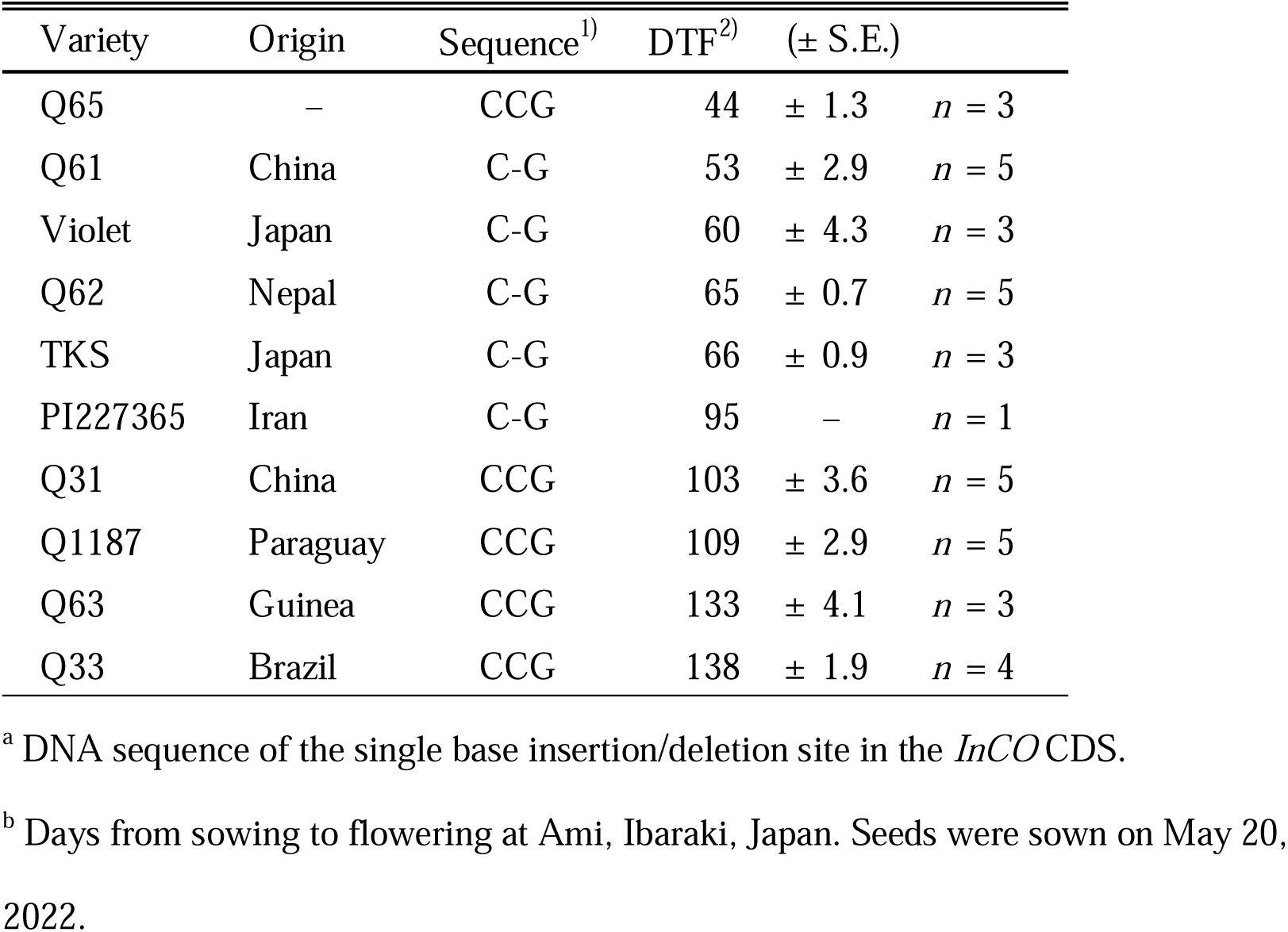
DTF and the single-base insertion/deletion in the CDS of *InCO*/*IhCO* among morning glory varieties.

| Variety | Origin | Sequence<sup>1)</sup> | DTF<sup>2)</sup> | (± S.E.) | |
| --- | --- | --- | --- | --- | --- |
| Q65 | – | CCG | 44 | ± 1.3 | $n = 3$ |
| Q61 | China | C-G | 53 | ± 2.9 | $n = 5$ |
| Violet | Japan | C-G | 60 | ± 4.3 | $n = 3$ |
| Q62 | Nepal | C-G | 65 | ± 0.7 | $n = 5$ |
| TKS | Japan | C-G | 66 | ± 0.9 | $n = 3$ |
| PI227365 | Iran | C-G | 95 | – | $n = 1$ |
| Q31 | China | CCG | 103 | ± 3.6 | $n = 5$ |
| Q1187 | Paraguay | CCG | 109 | ± 2.9 | $n = 5$ |
| Q63 | Guinea | CCG | 133 | ± 4.1 | $n = 3$ |
| Q33 | Brazil | CCG | 138 | ± 1.9 | $n = 4$ |

<sup>a</sup> DNA sequence of the single base insertion/deletion site in the *InCO* CDS.
<sup>b</sup> Days from sowing to flowering at Ami, Ibaraki, Japan. Seeds were sown on May 20, 2022.

DTFs of the accessions presented in Table 2 were investigated in 2022. They were from 44 days of Q65 to 138 days of Q33. The accessions without the SBD in the CDS of *InCO*/*IhCO* tended to have longer DTFs than those with the SBD, except for Q65 (**Table 2**).

### Comparison of InCO*/*IhCO expression between TKS and Q65

The expression levels of three transcripts, *InCO* (ni), *InCO* (si), and *InCO* (li) were compared between TKS and Q65 in SD and LD conditions using qRT-PCR (**Fig. 7**). The relative expression levels of the Q65 allele tended to be higher than TKS in all three transcripts across all conditions. Especially, the difference between TKS and Q65 was the most pronounced in *InCO* (si) (**Fig. 7B**). The expression level of *InCO* (si) was more than sevenfold higher in Q65 than in TKS after 14 h of darkness in the SD condition.

In all conditions, among the transcripts of three types, *InCO* (li) exhibited the highest expression level, as reported earlier in the relevant literature (**Liu et al. 2001**), followed by *InCO* (si). The lowest expression level was observed for *InCO* (ni) (**Fig. 7B, C**). Although quantities differed among the three transcripts, the expression patterns responding to the day length and the dark period were mutually similar. These tendencies were observed for both TKS and Q65.

When comparing expression levels of the transcripts encoding the functional InCO protein between TKS and Q65, the expression level of *InCO* (si) in Q65 was 52 times and 13 times, respectively, more than those of *InCO* (ni) in TKS at 8 h and 14 h after dark in SD conditions.

## Discussion

### InCO/IhCO, a candidate gene for the most significant QTL, qIF3, for DTF

In this study, four QTLs for DTF, *qIF1–4*, were detected in the F_2_ population of Q65 and TKS (**Table 1**). Among these QTLs, *qIF3* was the most significant QTL for DTF. It was detected in both 2011 and 2012. *InCO*/*IhCO* is located near the LOD peak of *qIF3*, and *InCO* is known to the ortholog of *CO* (**Liu *et al*. 2001**), which is a central regulator of photoperiodism in *Arabidopsis* (**Andres and Coupland 2012**). Therefore, *InCO*/*IhCO* is probably a candidate gene for *qIF3*. If *InCO* were *qIF3*, then *InCO*/*IhCO* would be expected to have polymorphisms causing functional differences between TKS and Q65. In fact, SBD causes a frame-shift in the *InCO*/*IhCO* CDS (**Fig. 5**). The expression analysis and splicing site prediction demonstrated that the transcript variant *InCO* (si) is the correctly spliced mRNA encoding the intact protein in the Q65 allele (**Fig. 6, Supplemental Fig. 6**). Actually, *InCO* (<u>ni)</u> was earlier thought to be a transcript without introns and encoding the functional InCO protein (**Liu *et al*. 2001**), but it appears to be an aberrant splicing transcript (**Fig. 6**, **Supplemental Fig. 6**). Although the splicing-variant *InCO* (ni) of TKS allele encodes functional CONSTANS-like protein (**Liu *et al*. 2001**), the expression level of TKS *InCO* (ni) is much less than that of Q65 *InCO* (si) (**Fig. 6**). Reportedly, the expression levels of *CONSTANS* affect the flowering time in *Arabidopsis* (**Rosas *et al*. 2014**). Therefore, the TKS allele of *InCO* might be defective because of the frame shift caused by the SBD. For that reason, we designate the TKS/Violet type defective allele as *inco-1* and the ancestral allele without the SBD as *InCO-2*.

### The allelic difference of InCO/IhCO corresponds with the effects of qIF3 on DTF

If *InCO*/*IhCO* were *qIF3*, then the effect of each *InCO***/***IhCO* allele on the DTF would be expected to match the effect of *qIF3*. The Q65 allele of *qIF3* has the effect of increasing DTF, whereas the TKS allele of *qIF3* has the effect of decreasing DTF (**Fig. 2**). Therefore, if *InCO***/***IhCO* were *qIF3*, then it can be inferred that *InCO*/*IhCO* allele without the SBD would increase DTF; also, *inco-1* would decrease DTF. This inference corresponds with results showing that the accessions bearing *inco-1* tend to flower earlier than those bearing the *InCO*/*IhCO* allele without the SBD, except Q65 (**Table 2**). Among the strains examined for this study, Q65 flowers the earliest, but it has the *IhCO* allele without the SBD (**Table 2**). The fact that Q65 flowers the earliest among accessions examined for this study does not contradict the notion that the *InCO*/*IhCO* allele without the SBD has an effect of delaying flowering. If *InCO***/***IhCO* is *qIF3*, then the Q65 allele, that is the *IhCO* allele without the SBD, would also exert a delaying effect on flowering because the Q65 allele of *qIF3* has the effect of increasing DTF (**Table 1**, **Fig. 2**). Consequently, the allelic differences in the structure and the predicted effect for DTF of *InCO***/***IhCO* can well explain the allelic differences of *qIF3*.

In rice, dysfunctional alleles of *Hd1* (*hd1*), the rice ortholog of *CONSTANS*, are known to reduce photoperiod sensitivity. Moreover, they have the effect of promoting heading in field conditions (**Ebana *et al*. 2011**, **Fujino *et al*. 2019**, **Hori *et al*. 2016**, **Mo *et al*. 2021, Yano *et al*. 2000**). The early flowering tendency of rice cultivars with *hd1* corresponds with that of *I. nil* varieties with *inco-1* (**Table 2**). The photosensitivity of crops with ancestors originating from tropical regions, which are classified as short-day plants, tend to be reduced or lost during domestication and expansion to high-latitude areas (**Lin *et al*. 2021**). The *I. nil* varieties with *inco-1* were found exclusively in Asia but not in the Americas, the origin of *I. nil*, and Africa. Consequently, *inco-1* might have emerged in Asia, and might have contributed to the spread of *I. nil* across temperate Asia by reducing its photoperiod sensitivity.

### Genetic mechanism for the early flowering of Q65

The fact that Q65 was the earliest flowering among the strains investigated for this study (**Table 2**) despite having the ancestral allele of *InCO***/***IhCO* implies that Q65 possesses a mechanism for early flowering that is distinct from that of Asian *I. nil* strains. Genes other than *IhCO* might be responsible for causing Q65 to flower early.

Four QTLs for DTF were detected in this study. Among the four QTLs, the Q65 allele of *qIF1* and *qIF4* have the effect of reducing DTF and enhancing flowering (**Table 1**, **Fig. 2**). However, the Q65 allele of two major QTLs, *qIF2* and *qIF3*, delayed flowering. The sum of effects of these QTLs was greater than those of *qIF1* and *qIF4* (**Table 1**, **Fig. 2**). These detected QTLs were inadequate to explain the early flowering of Q65 entirely. The method for QTL mapping used for this study cannot detect epistatic interactions among more than two QTLs. Therefore, additional genetic studies using suitable experimental lines might be necessary to elucidate the causal genetic mechanism of the early flowering in Q65.

In rice, the combination of the loss of function of two genes, *Ghd7* and *Osppr37*, results in extremely early flowering. Given this genetic background, *Hd1* has been reported to promote earlier heading (**Fujino *et al*. 2018)**. Similarly, in Q65, the loss of function of multiple genes other than *IhCO* can be presumed to contribute to early flowering.

### Polymorphism in the 5***′***-upstream regulatory region of InCO*/*IhCO

Insertion of a SINE-like retrotransposon, which does not exist in TKS allele of *InCO***/***IhCO*, was observed in 5′-flanking region of Q65 allele (**Fig. 4**). SINEs are known to cause changes in the chromatin structure and to affect gene expression (**Elbarbary *et al*. 2016**). The expression levels of Q65 allele of *InCO***/***IhCO* are greater than those of TKS (**Fig. 6**). Therefore, structural differences and the SINE-like sequence insertion upstream of Q65 allele might affect the expression level of *InCO***/***IhCO* (**Supplemental Fig. 5**).

### An intron-containing InCO/IhCO transcript variant, InCO *(li)*

In both Q65 and TKS, *InCO* (li) exhibits the highest expression level among the three splicing variants, although it includes an intron sequence and encodes truncated protein without CCT domain (**Liu *et al*. 2001**) (**Fig. 7A**). In *Arabidopsis,* an intron-containing *CO* transcript variant, *CO*β, also encodes a truncated protein without CCT domain caused by premature stop codon. The COβ protein interacts with the functional form of CO protein, COα, and reduces its stability (**Gil *et al*. 2017**). Therefore, the InCO (li) protein might also interact with the functional InCO/IhCO protein and might affect *InCO*/*IhCO* functions. However the expression level of *CO*β is less than *CO*α (**Gil *et al*. 2017**). Therefore, the highest expression level of *InCO* (li) among splicing variants is a unique and interesting feature of *InCO*/*IhCO*.

## Conclusion

This study suggests that the *CONSTANS* ortholog of morning glory, *InCO*/*IhCO*, similarly to the *Hd1* gene in rice, suppresses flowering in the long-day condition and suggests that it is involved in photoperiodic flowering. Consequently, both dicot and monocot short-day plants might share a common mechanism mediated by *CONSTANS* orthologous genes that suppress flowering under long-day conditions. In addition, this study suggests that the SBD which occurred in the CDS of *InCO* might have contributed to the adaptation of *I. nil*, originally native to the tropical regions of the Americas, to temperate Asia.

### Author Contribution Statement

Conceptualization: HK and TK; resources provision: EN and AH; mapping population development: EN; DNA marker development: TI, HK, HF, SS, and SI; investigation: HK, TI, KE, and TK; writing – original draft preparation: HK and TK; writing – review and editing: TK, EH, and AH; supervision: AH, EN, MO, and NW; project administration: TK; funding acquisition: TK and MO. All authors have read and agreed to the published version of the manuscript.

## Acknowledgments

This work was partially supported by JSPS KAKENHI Grant Number 18K05569. This work was also partially supported by the Cooperative Research Grant of the Plant Transgenic Design Initiative (PtraD) at the Gene Research Center, University of Tsukuba.

