## Supplemental Table 1 for "Frame-shift mutation of *InCO* might cause early flowering of Japanese morning glory and might have contributed to northward expansion"

**Supplemental Table 1.** List of EST-SSR markers derived from *I. batatas*

| Marker name | Core motifs | Chromosome | Forward primer (5´ to 3´) | Reverse primer (5´ to 3´) |
|---|---|---|---|---|
| IES1076 | AAC | 2 | TAGAGCCTTTGACGTCCGAT | TAATTTTCTCCAATTGCCGC |
| IES0411 | ATC | 3 | AGGAAGCCCAATGGAGTTTT | AGTCTGGGTTGGCAATTACG |
| IES0856 | GGA | 4 | TTGCTGATTCGGACACTGAG | CACTCCCAAGAAGTTGCTCC |
| IES0575 | AG | 4 | GCTGATAACACCTGTGAGAGATAGA | TGTGTTCAATTTCACCGGAA |
| IES0160 | ATC | 5 | AACGGCCAGAGATTGTGAAG | CATGGCAAATCATCGTCATC |
| IES0847 | AAG | 5 | CGGATCACTGAGATGAGGGT | TGTGAGCCATTCTGCGTAAC |
| IES0809 | ATC | 8 | AAGCACCTACTCCTGCTCCA | TCCACCCTGCTCTAGGTACG |
| IES0902 | AAG | 9 | CGTAGCTCTCCCCTCTCCTT | TGCTTGTTGCTGTTTCAACC |
| IES0410 | AAG | 10 | CAGGGGCATAACCGTAATTG | ATGATCCCCAGGTTTAAGGC |
| IES0433 | ATC | 12 | CCCATTTCTCCACAGGGATA | GAAGGGCAGGAACAAATTCA |
| IES0874 | ATC | 12 | AGATGCACTCCCTTGCTTTC | ATGGGGAGGAGATCCAAGTT |
| IES0414 | AG | 12 | TGCATTTCTCTCTGTTCCCC | GTAAAAATCCCCGTGCTGAA |
| IES0315 | GGC | 13 | CTATAGGTTGGCACCGGAAA | TTGTGGGCTATATTGGCTCC |
| IES0680 | AAG | 13 | GGGTTTTCAGAGACCGTTGA | CCATGCACACCTTCACAAAC |
| IES0536 | AAG | 13 | ATGTTGCCTAATGGACAGGC | AATGGGACAAATGGCAACTC |
| IES0862 | ATC | 14 | TCGTTGGGACTCCTGATACC | AGTTGTCGCTTGCGTTTCTT |
| IES0177 | GGA | 15 | CAAACATGGACGAAATCACG | TCACCGTCGTCATCGTACTC |
