## Supplemental Table 2 for "Frame-shift mutation of *InCO* might cause early flowering of Japanese morning glory and might have contributed to northward expansion"

**Supplemental Table 2.** HRM-SNP and SSR markers developed and used for genetic linkage-map construction and QTL analysis of DTF in *I. nil* and *I. hederacea*

| Marker name | Marker type [a)] | Chromosome | Forward primer (5′ to 3′) | Reverse primer (5′ to 3′) |
|---|---|---|---|---|
| rJMSF039K04.591 | HRM | 1 | TCTTCCTGTGCAACTTACGC | AAATTGCTCCATTTGGCTTG |
| Contig781.719 | HRM | 1 | AACTCTTGCTCGAGCCATTT | CACTCAGGTTCCTGATTTTCTAA |
| fe-SSR | SSR | 2 | TCAAACATCGACCTCGTGAC | GCCCTCCATCTCTCTTCACA |
| Contig8667.442-m | HRM | 3 | CGTGTCCTTGTCGTTTTCATT | CCCTTCAGCTCTTCCCTCTT |
| MYBX | HRM | 3 | GCCATCGCCAACATTTTAC | TTGATCTAGATTGTGGGCAAGA |
| Contig1617.421-dy | HRM | 3 | CCATGGACAAAACCCAGAAC | CGGACTGATGGTGCTGTAGA |
| Contig1620.744 | HRM | 3 | CACGACCAAACAGCTCTTCA | CGTACAGATCGACCTGATTGC |
| PnFT2 | HRM | 3 | ATCAAATAGGTTGCCGGAGA | CGACGACCAGTGGGTCTCT |
| Contig10056.454 | HRM | 4 | CTCCAATCATCTTCAACACCAA | CACGACATGATCGTTCCTCA |
| InPHYC | HRM | 4 | CTGCACTTGAAAAAGCAGCA | GCTTACCCGAATTTTTGCAG |
| Contig2109.295-mg | HRM | 4 | GCCGCTCAGTTCTTGAAAGT | GCGAAGACGAGGTCTTGATAG |
| Contig5247.271-a3 | HRM | 5 | AAAGCCGACACGAATTTGAC | TGGCTTCATCAAAGCTTCCT |
| InMYB2 | HRM | 5 | ATGGTTACATGTGTTTAGGTGGTC | CTTCACATCGTTCGCTGTTC |
| Contig12385.595 | HRM | 5 | ATCTTCATCGGTGACGCTTT | GTGGTCCCCGGAGACTTT |
| Contig7414.299 | HRM | 5 | AACACCATGGGAGATTCGAG | GGCAAGATCGATTTCTTCGT |
| Contig6884.391 | HRM | 5 | TGGTGGCACAATAATTGACATT | AGCCAGCACAGTGTTTTTGA |
| Contig126.513 | HRM | 5 | GTCGCTTCGCAAATCCTAGA | TATCTGATGGCCAAGGGTTG |
| Contig11461.715 | HRM | 5 | CGACAATCCAAAATCTGCAA | ACCTGCACACTGAAGCTCAC |
| Contig7935.1121 | HRM | 5 | CTGCTGTAAGGCCTCCACCT | AAGGGAGATTCGCGCTAAAG |
| Contig1107.272 | HRM | 5 | CAGATGCAGCCCAATTGATA | TCCAATTTCAATGGTCCTTTT |
| InG31484728_Ca | HRM | 5 | GGAGATTACAGTTCCGACGTC | TTTGTCCTCCTCCGTCGAC |
| Contig10004.219-efp | HRM | 5 | TTTCTCCAAATAAAGTACACTGCAA | GACACAATTTCTGGGTCCAA |
| DP-SSR | SSR | 6 | ATGGTGGCATGGACAATCTT | CCTTCACCCATGTAGTTCCTG |
| InPHYE | HRM | 6 | GGCCATAGCTCAGTACAATGC | TCTTCTTCAGTCACATTCTGTGG |
| Contig10633.676-ivs | HRM | 7 | GCCTTGTCCATTTTCGTCAC | TCAACGAGCGGTTCATTATC |
| Contig12273.307-kbt | HRM | 7 | CAGTCCACCTCTGGGAATCT | CGGTGAATGTATGGGCTGAT |
| JMFN043J13.0370 | HRM | 9 | GCGTTTGAACTCCAGATTGC | CAAAACTCATCGGCAAACAA |
| PnFT1 | HRM | 9 | CCGGCAGACAGTTTATGCAC | TCAGCAAAGTTTCGAGTGTTG |
| PnTFL1a | SSR | 10 | GAGAACAAGAAGCCCCTCAA | ACTGTGGGAGTGAAGGCATC |
| Contig1647.128-pr | HRM | 10 | GAGAGAGTCACGTTAATCCTGAGA | GTGGGGGTAGTGTTGAAGGA |
| Contig3344.254 | HRM | 10 | GCTCCTCACACCTGTCATTT | TCTCCCCTTCTTCATCTCCA |
| Contig4567.156-InCO | HRM | 11 | CAGAAGTGGCAACAAGCAGA | GAACGGCATATGTCGCAGA |
| Contig4216.592-InGI | HRM | 11 | GGCATGCTGTTATTCATCCA | GGCAAAAGGTGGACTATTTTT |
| JMFF032K01.306-c1 | HRM | 11 | TTGCAGATTGAGATGGTTGAA | GCTTATGGAGGCGTAGAATGA |
| Contig2647.823-sp | HRM | 11 | TGATATCTGTTTTATTTGGGAGTGG | CAAAATAAAAACAAAAGTAATCCGTA |
| rJMFF001I14.223 | HRM | 12 | GTGTATGGAGCTGGCTTGG | GCACGCTCTCTCAAAGATGA |
| Contig2300.318 | HRM | 12 | AACAGCATCCGAGTCTGGAA | CCCGTGGCTACGAGATAGTG |
| Contig3548.340 | HRM | 12 | AAGCTAAGGTGTGGGAGAAGC | GTGGCATTGGTTTAGGCATT |
| Contig12.373 | HRM | 12 | GGGGATTGTGATCATTCTGG | TGCCAATCTCGTAATCCTCA |
| Contig23.333-r3 | HRM | 13 | ACCTGGTGAACCACGGGATA | CTGACCGAAAAACCCCTCTC |
| CHS-E | HRM | 14 | AGCGATCCAGAAAAGGGAGT | TCCTGGGCTCATATCAAAGC |
| JMFS127D02.495 | HRM | 14 | ACGGATTTGAGGACCCTTTT | AAATTGGGTTGAGCAAGGTG |

[a)] HRM and SSR respectively represent high-resolution melt and simple sequence repeats.
