## Supplemental Table 3 for "Frame-shift mutation of *InCO* might cause early flowering of Japanese morning glory and might have contributed to northward expansion"

**Supplemental Table 3.** List of developed Tm-shift primers based on the *I. nil* EST database

| Marker name | Chromosome | Fwd. / Rev. | Primer sequences (5´ to 3´) |
|---|---|---|---|
| Contig71.0517 | 1 | Forward | AAGAAGCTTTCGAGGCGGCGTA |
| | | Reverse 1 | GCGGGCAGGGCGGCGCGGCGGAATCACTGGTCATT |
| | | Reverse 2 | GCGGGCGCGGCGGAATCACTGGTCATA |
| Contig73.0355 | 1 | Forward | GAAGCAGAATGTTAAGCAATCGAGGA |
| | | Reverse 1 | GCGGGCAGGGCGGCTATTCAGAGTAACAGGAGTAGGCGCG |
| | | Reverse 2 | GCGGGCTATTCAGAGTAACAGGAGTAGGCTCA |
| Contig213.0351 | 2 | Forward 1 | GCGGGCAGGGCGGCCGGCGAATTTGGTGACGTATCTC |
| | | Forward 2 | GCGGGCCGGCGAATTTGGTGACGTATCTG |
| | | Reverse | CGAGTTGCTCATCTTCGCCACAT |
| Contig13313.0335 | 2 | Forward | TGTGCAATAGCTTTATTCGGAACCA |
| | | Reverse 1 | GCGGGCAGGGCGGCGGAGAGCAAGAATTTCCATGACCGG |
| | | Reverse 2 | GCGGGCGGAGAGCAAGAATTTCCATGACCGC |
| Contig29.0586 | 2 | Forward | TGTTTCACGCATGGGAAGTATGAGAA |
| | | Reverse 1 | GCGGGCAGGGCGGCAACTGCACTTAAGTTGGCAGGGTCCG |
| | | Reverse 2 | GCGGGCAACTGCACTTAAGTTGGCAGGGTACA |
| Contig117.0507 | 2 | Forward | CGGGTGTTCTTCAACAGCTCGAC |
| | | Reverse 1 | GCGGGCAGGGCGGCCACGAGTCGGTGCCAAGACAAG |
| | | Reverse 2 | GCGGGCCACGAGTCGGTGCCAAGACCAA |
| Contig39.0484 | 2 | Forward 1 | GCGGGCAGGGCGGCGCTTCATCGTTTCTTACCTCGTGG |
| | | Forward 2 | GCGGGCGCTTCATCGTTTCTTACCTCGGGA |
| | | Reverse | GTAATGCCCACGAACAGCAGAAGC |
| Contig341.0578 | 3 | Forward | GCGAGATGATTTGGATAACATTTCCCT |
| | | Reverse 1 | GCGGGCAGGGCGGCAGAAGTTCTTCTCAAAGCGAACCCAG |
| | | Reverse 2 | GCGGGCAGAAGTTCTTCTCAAAGCGAACCAAA |
| Contig591.0478 | 3 | Forward | GGGAAGCATTCCTGCAAAGGTTTAGA |
| | | Reverse 1 | GCGGGCAGGGCGGCGCGAGTCTCCCACAAACATAATCGTC |
| | | Reverse 2 | GCGGGCGCGAGTCTCCCACAAACATAATCTTT |
| Contig11523.0689 | 3 | Forward 1 | GCGGGCAGGGCGGCCTAAACCTTCCTTGGTTGTGAGGCGG |
| | | Forward 2 | GCGGGCCTAAACCTTCCTTGGTTGTGAGGAGA |
| | | Reverse | ACAACCGGAAGGTCAAGAAAATCTGG |
| Contig132.0440 | 4 | Forward 1 | GCGGGCAGGGCGGCGCGCAGTCGTACATGGATGACTTG |
| | | Forward 2 | GCGGGCGCGCAGTCGTACATGGATGACGTA |
| | | Reverse | AATGAAACTGCCAAGAACCAGGGC |
| Contig210.0518 | 4 | Forward | CGAAGGTGGCAAAATATATGGGGTG |
| | | Reverse 1 | GCGGGCAGGGCGGCCAATTGCAAAAGGCTGCTCCAAGAC |
| | | Reverse 2 | GCGGGCCAATTGCAAAAGGCTGCTCCAATAT |
| Contig10534.0459 | 4 | Forward | GATACTGAGGTCGGTGGGAATCAAGG |
| | | Reverse 1 | GCGGGCAGGGCGGCTGCTGATACTAAGGTCCCTGGTATGG |
| | | Reverse 2 | GCGGGCTGCTGATACTAAGGTCCCTGGTAGGA |
| Contig618.0053 | 4 | Forward | CGCCCCTCTCCTCCTCCTCTTA |
| | | Reverse 1 | GCGGGCAGGGCGGCCATGGAAACGGCGAACTGG |
| | | Reverse 2 | GCGGGCCATGGAAACGGCGAACGGA |
| Contig312.0049 | 5 | Forward 1 | GCGGGCAGGGCGGCACATTCTCCGCCGTTTTCTGAGTGTC |
| | | Forward 2 | GCGGGCACATTCTCCGCCGTTTTCTGAGTTTT |
| | | Reverse | ATTGGGGATTGGATTGTGATGTATGC |
| Contig9549.0185 | 5 | Forward 1 | GCGGGCAGGGCGGCCATTCATGGCAGCTTATGCAACAGC |
| | | Forward 2 | GCGGGCCATTCATGGCAGCTTATGCAACAGG |
| | | Reverse | CGAGAACCTTCTCCAGAGCCACTTC |
| Contig11034.0239 | 5 | Forward | TGCATATGCTACAGGAATTTCTCCCA |
| | | Reverse 1 | GCGGGCAGGGCGGCCGTTTGATGGTCATACCATGCATTCC |
| | | Reverse 2 | GCGGGCCGTTTGATGGTCATACCATGCATGCT |
| Contig10043.1070 | 5 | Forward | GTCGGTAATGAGGTTCTGGCGAAGAG |
| | | Reverse 1 | GCGGGCAGGGCGGCTTTCCCACCTCTCCAATCTCAACCAG |
| | | Reverse 2 | GCGGGCTTTCCCACCTCTCCAATCTCAACAAA |
| Contig223.0210 | 5 | Forward 1 | GCGGGCAGGGCGGCATGCGAGTTGTTTGAATTGGACTGG |
| | | Forward 2 | GCGGGCATGCGAGTTGTTTGAATTGGACGGA |
| | | Reverse | TGATCGAAAATCAATAGCACACAAA |

**Supplemental Table 3.** (continued)

| Marker name | Chromosome | Fwd. / Rev. | Primer sequences (5´ to 3´) |
|---|---|---|---|
| Contig1732.1500 | 6 | Forward | GACCTGCCGGAGAAGGTTTTGTTAGA |
| | | Reverse 1 | GCGGGCAGGGCGGCACACATTCTCTCTGTGTCCAAAGTCG |
| | | Reverse 2 | GCGGGCACACATTCTCTCTGTGTCCAAAGGCA |
| Contig348.0441 | 6 | Forward | GCCGCAGCCTCGAATTGTTAGT |
| | | Reverse 1 | GCGGGCAGGGCGGCGGCGCAATGAGTGGTAAAATCG |
| | | Reverse 2 | GCGGGCGGCGCAATGAGTGGTAAAAGCA |
| Contig638.0218 | 6 | Forward | CGATGTGCCGGTGATTGACCTAC |
| | | Reverse 1 | GCGGGCAGGGCGGCCCGAATCTCCCCACTCCTCAGAG |
| | | Reverse 2 | GCGGGCCCGAATCTCCCCACTCCTCAGAC |
| Contig10207.0412 | 6 | Forward 1 | GCGGGCAGGGCGGCCCATGTGTTCTCGCTCCCAAATAGG |
| | | Forward 2 | GCGGGCCCATGTGTTCTCGCTCCCAAATAGC |
| | | Reverse | AAGCAGAAAATCCCATGTCCAATCG |
| Contig13203.0679 | 7 | Forward 1 | GCGGGCAGGGCGGCGCTGTTGACGCTTCAAAATGCTTGTC |
| | | Forward 2 | GCGGGCGCTGTTGACGCTTCAAAATGCTTTTT |
| | | Reverse | ACTGATGAGAGAAGCACGGACAGCTT |
| Contig7688.0677 | 7 | Forward 1 | GCGGGCAGGGCGGCGCACTGCCTAGCTCAGCCTCCTCTA |
| | | Forward 2 | GCGGGCGCACTGCCTAGCTCAGCCTCCTCTT |
| | | Reverse | TTGTAGGTGCCAATACCAATGCCAA |
| Contig361.0384 | 7 | Forward | TTCGCTACTTCATCCACCAGTTTTGG |
| | | Reverse 1 | GCGGGCAGGGCGGCCAGCAGAAGAACCAAAAGAAAATTCG |
| | | Reverse 2 | GCGGGCCAGCAGAAGAACCAAAAGAAAATGCA |
| Contig13068.0510 | 7 | Forward 1 | GCGGGCAAGTTGCTCCAAGAAGAACAGGGAGA |
| | | Forward 2 | GCGGGCAGGGCGGCAAGTTGCTCCAAGAAGAACAGGGCGG |
| | | Reverse | CTCCACAAAGCATTCCCTTCACAGAC |
| Contig45.0110 | 8 | Forward 1 | GCGGGCAGGGCGGCAAACCGGCAAAGAAAGGTGGGGAC |
| | | Forward 2 | GCGGGCAAACCGGCAAAGAAAGGTGGGGAG |
| | | Reverse | TTTGACGGGAAATCATCGGAAAAA |
| Contig12633.0348 | 8 | Forward 1 | GCGGGCAGGGCGGCCTTGATATGGCTTGCCGATGAACGTC |
| | | Forward 2 | GCGGGCCTTGATATGGCTTGCCGATGAACTTT |
| | | Reverse | GCAGAAGGATCTGTTGGAGACAATGG |
| Contig14002.0191 | 8 | Forward | TCAAACCCTCCAGAACCTCCCTTTT |
| | | Reverse 1 | GCGGGCAGGGCGGCTGGTGATGAAGAAGATTGCTCGTCG |
| | | Reverse 2 | GCGGGCTGGTGATGAAGAAGATTGCTCGTCC |
| Contig6146.0392 | 9 | Forward 1 | GCGGGCAGGGCGGCCACCCTCGATAATTCCTTCCAACTCA |
| | | Forward 2 | GCGGGCCACCCTCGATAATTCCTTCCAACTCT |
| | | Reverse | AAAATACCCGATTCCGCATGAAATTG |
| Contig502.0119 | 9 | Forward | CCAGTGATTGATTGATCTCTCGTTACCA |
| | | Reverse 1 | GCGGGCAGGGCGGCTCTGCATCTTGCCTCTCTTTCTCGTC |
| | | Reverse 2 | GCGGGCTCTGCATCTTGCCTCTCTTTCTCTTT |
| Contig572.0192 | 9 | Forward | AGGGGGAGAAGATGAAGGAGAGAACC |
| | | Reverse 1 | GCGGGCAGGGCGGCACTTCCACTGTGTGTAGCGCTCGGTC |
| | | Reverse 2 | GCGGGCACTTCCACTGTGTGTAGCGCTCGTTA |
| Contig1098.0735 | 9 | Forward | TCCGTTCACGCATCCTTCCACT |
| | | Reverse 1 | GCGGGCAGGGCGGCTGACCCGGTTCTTCAGATCAGG |
| | | Reverse 2 | GCGGGCTGACCCGGTTCTTCAGATCCGA |
| Contig31.0114 | 10 | Forward | AGAAACCAAAACGGAACAAATACGCC |
| | | Reverse 1 | GCGGGCAGGGCGGCAAGTCATGGAAGCTAAGAGAGAGAAG |
| | | Reverse 2 | GCGGGCAAGTCATGGAAGCTAAGAGAGAGCAT |
| Contig9165.0662 | 10 | Forward 1 | GCGGGCAGGGCGGCTGCATTCTTTGGGAAAACTTGGTCC |
| | | Forward 2 | GCGGGCTGCATTCTTTGGGAAAACTTGGTCG |
| | | Reverse | TAGTGGTTGGAACTCACGCTGTGCT |
| Contig52.0536 | 10 | Forward | AAAAGTGGGCTCTTTTGTTGACAGGC |
| | | Reverse 1 | GCGGGCAGGGCGGCCAAAAAACACATGCAATCCCATTGCG |
| | | Reverse 2 | GCGGGCCAAAAAACACATGCAATCCCATTTCA |
| Contig683.0110 | 10 | Forward | GCCACAGTGACACTCCAATGTTGA |
| | | Reverse 1 | GCGGGCAGGGCGGCCCGGCCAAAGGTCCGGTAAAGTA |
| | | Reverse 2 | GCGGGCCCGGCCAAAGGTCCGGTAAAGTT |

**Supplemental Table 3.** (continued)

| Marker name | Chromosome | Fwd. / Rev. | Primer sequences (5′ to 3′) |
|---|---|---|---|
| Contig514.0130 | 10 | Forward 2 | GCGGGCACGCCCCACAGTCAACGACA |
| | | Forward 1 | GCGGGCAGGGCGGCACGCCCCACAGTCAACGCCG |
| | | Reverse | GGCAACTCGGTGAGTGAGGCTT |
| Contig13171.0174 | 11 | Forward | AATCTGGGGAGTTGACCCAAAAAGC |
| | | Reverse 1 | GCGGGCAGGGCGGCCTTGGTCTCCTTGATCTACGGCTCG |
| | | Reverse 2 | GCGGGCCTTGGTCTCCTTGATCTACGGCTCC |
| Contig394.0210 | 11 | Forward | TCTGGCCAAGGATAAAGCGTTT |
| | | Reverse 1 | GCGGGCAGGGCGGCATATCGGCGGCGAGGTGTGTC |
| | | Reverse 2 | GCGGGCATATCGGCGGCGAGGTGTGTG |
| Contig10076.0370 | 11 | Forward | CCACATTCTCTAGCTGCTCGATTGCT |
| | | Reverse 1 | GCGGGCAGGGCGGCAGTACTTCGACATGCCACCTTTGCCG |
| | | Reverse 2 | GCGGGCAGTACTTCGACATGCCACCTTTGACT |
| Contig529.0215 | 11 | Forward 1 | GCGGGCAGGGCGGCGTTCCGAAGCGGTGGACAGTTGAG |
| | | Forward 2 | GCGGGCGTTCCGAAGCGGTGGACAGTTTAA |
| | | Reverse | TCTTCTTCGGCGCACTCATTTCTC |
| Contig43.0622 | 11 | Forward | GCAACAGTCCATGAAGAAAGAGCGAA |
| | | Reverse 1 | GCGGGCAGGGCGGCTCTTAGTATCCACGGCGGTGGTACTG |
| | | Reverse 2 | GCGGGCTCTTAGTATCCACGGCGGTGGTAATA |
| Contig639.0355 | 12 | Forward 1 | GCGGGCAGGGCGGCGGGACCTAATGGTGCCGAATGCC |
| | | Forward 2 | GCGGGCGGGACCTAATGGTGCCGAATTCA |
| | | Reverse | ATCTCACCGGCTTGGCGTACAAC |
| Contig50.0300 | 12 | Forward | TTCTCTGCGCAATCAAAGCCGTA |
| | | Reverse 1 | GCGGGCAGGGCGGCTGGATTGGTTGGTCCAAGATTCC |
| | | Reverse 2 | GCGGGCTGGATTGGTTGGTCCAAGATGCT |
| Contig268.0442 | 12 | Forward 1 | GCGGGCAGGGCGGCCGGATATGATTGATGGATTCCAGCAG |
| | | Forward 2 | GCGGGCCGGATATGATTGATGGATTCCAGAAA |
| | | Reverse | GTAGATGATCGGAATACCGAGCCGGG |
| Contig8337.0661 | 12 | Forward 1 | GCGGGCAGGGCGGCCGGCGAACGTAGCGTATAAACTCCG |
| | | Forward 2 | GCGGGCCGGCGAACGTAGCGTATAAACTCCC |
| | | Reverse | CGCCAACTTCACCAAGTACTTCACC |
| Contig114.0547 | 14 | Forward | GCCAATGTTTCCAAGGCTTCATGTT |
| | | Reverse 1 | GCGGGCAGGGCGGCGAGGAGCTCTTGGTCCTCCCTTGTC |
| | | Reverse 2 | GCGGGCGAGGAGCTCTTGGTCCTCCCTTTTT |
| Contig136.0560 | 14 | Forward 1 | GCGGGCAGGGCGGCCGATCGTTCCGTTTCTGCAAGC |
| | | Forward 2 | GCGGGCCGATCGTTCCGTTTCTGCAAGG |
| | | Reverse | ATTCCGACTTCATTGCAGGGCA |
| Contig539.0530 | 14 | Forward | GGTCTGAATAACGTTGTGTCGAAAA |
| | | Reverse 1 | GCGGGCAGGGCGGCAGATGCTGAGAAAACCTTGTGCAAG |
| | | Reverse 2 | GCGGGCAGATGCTGAGAAAACCTTGTGCCAA |
| Contig13.0186 | 15 | Forward 1 | GCGGGCAGGGCGGCCGGGAGAAGATTCTTAAGGACACG |
| | | Forward 2 | GCGGGCCGGGAGAAGATTCTTAAGGACCCA |
| | | Reverse | CGAACGATACAAAGGCAGCTGGAA |
| Contig6233.0433 | 15 | Forward | CCTCGCACTCCAACCTGCTTTACAT |
| | | Reverse 1 | GCGGGCAGGGCGGCGTCGAAGGAAAACGAAAACGTCGTC |
| | | Reverse 2 | GCGGGCGTCGAAGGAAAACGAAAACGTCGTG |
