## Supplemental Table 4 for "Frame-shift mutation of *InCO* might cause early flowering of Japanese morning glory and might have contributed to northward expansion"

**Supplemental Table 4.** Primers used for PCR amplification and DNA sequencing of *IhCO*

| Primer name | Sequences (5′ to 3′) | Fwd / Rev | Position[a] |
|---|---|---|---|
| InCO_pro2F | ACGGTTGTAGCAGTGCAGAACGAAAGAA | Forward | 7,614,486 - 7,614,513 |
| InCO_pro4F | CTTGTGGCTGGCTGTGATTCTG | Reverse | 7,616,393 - 7,616,414 |
| InCO_proQ65F | GATGCAATTGAGTTGATTAGCC | Forward | – |
| InCO_proQ65_5F | CGTTGGACGAAGAAGCCCT | Forward | – |
| InCO_proQ65R | CGAAGTGTGAGTCCTAGTATGAACC | Reverse | – |
| InCOpro3R | AAGGTAGGGGCAGTCATAGC | Reverse | – |
| InCO_midF | AAGCCATTGACAGGACAGGT | Forward | 7,616,327 - 7,616,346 |
| InCO_pro2R | ATCTTGTGGCTGGCTGTGATTCTGGTGG | Reverse | 7,616,389 - 7,616,416 |
| InCO_TerF | CATGATGGTTGGGCTGACGG | Forward | 7,617,026 - 7,617,045 |
| InCO_midR | GCATCCTGAGTCAAGAACCC | Reverse | 7,617,069 - 7,617,088 |
| InCO_Ter2F | AGTCCTGTTTCTCCTTCGGG | Forward | 7,617,673 - 7,617,692 |
| PnCO-realR | CTTTCGAGGCTCTTGAGTGG | Reverse | 7,617,855 - 7,617,874 |
| InCO_Ter2R | CGATAGCTGGAATAGTTAATGGGC | Reverse | 7,618,263 - 7,618,286 |

[a] Positions in the *I. nil* genome (Asagao 1.2).
