## Supplemental Fig. 1 for "Frame-shift mutation of *InCO* might cause early flowering of Japanese morning glory and might have contributed to northward expansion"

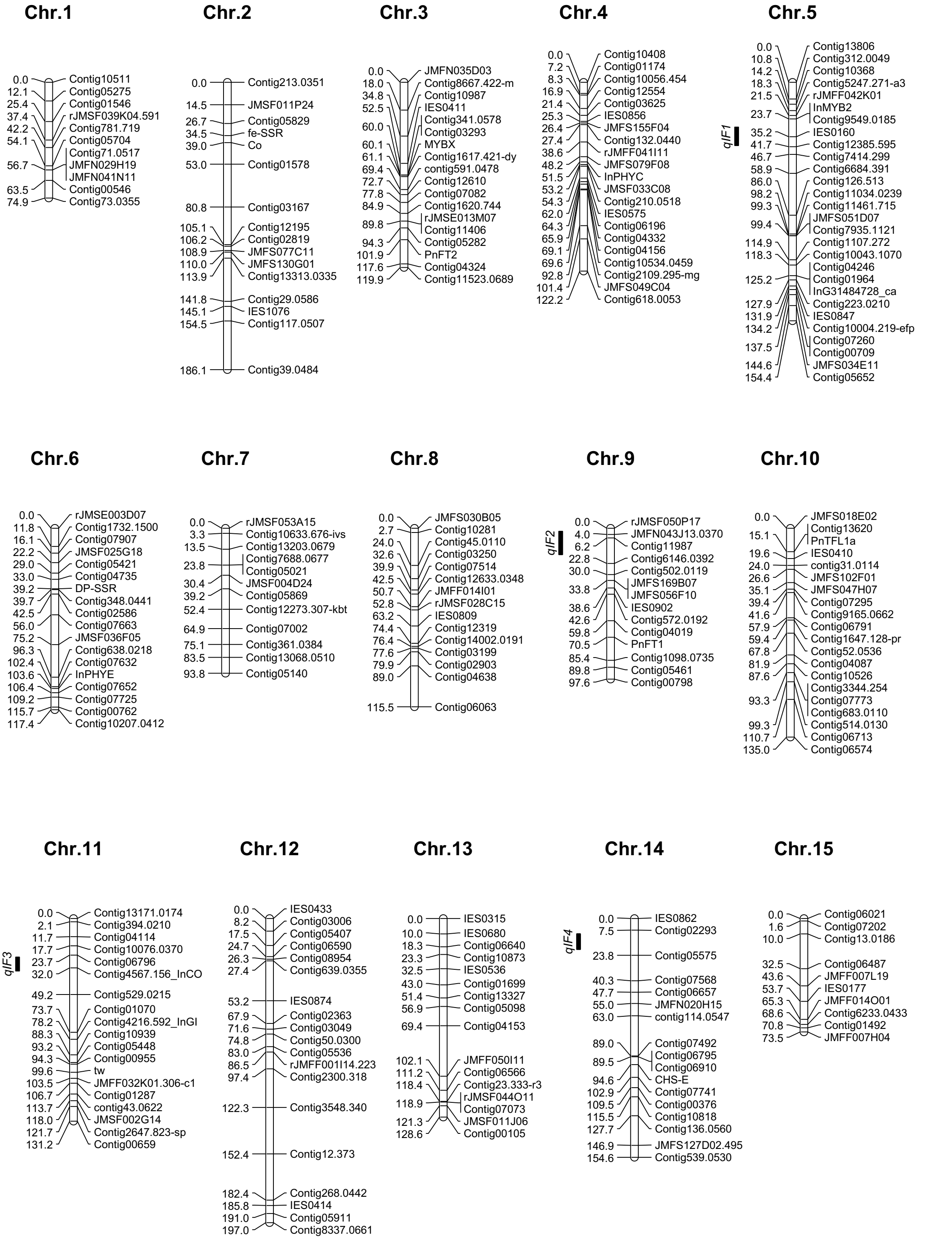

**Supplemental Fig. 1.** Genetic linkage map of $F_2$ population derived from the cross between Q65 × TKS and cultivated in 2011. The marker distance (cM) is presented on the left side of each chromosome. Marker names are shown on the right side. Vertical black bars show QTL regions for flowering days with 99% creditable intervals.
