## Supplemental Fig. 2 for "Frame-shift mutation of *InCO* might cause early flowering of Japanese morning glory and might have contributed to northward expansion"

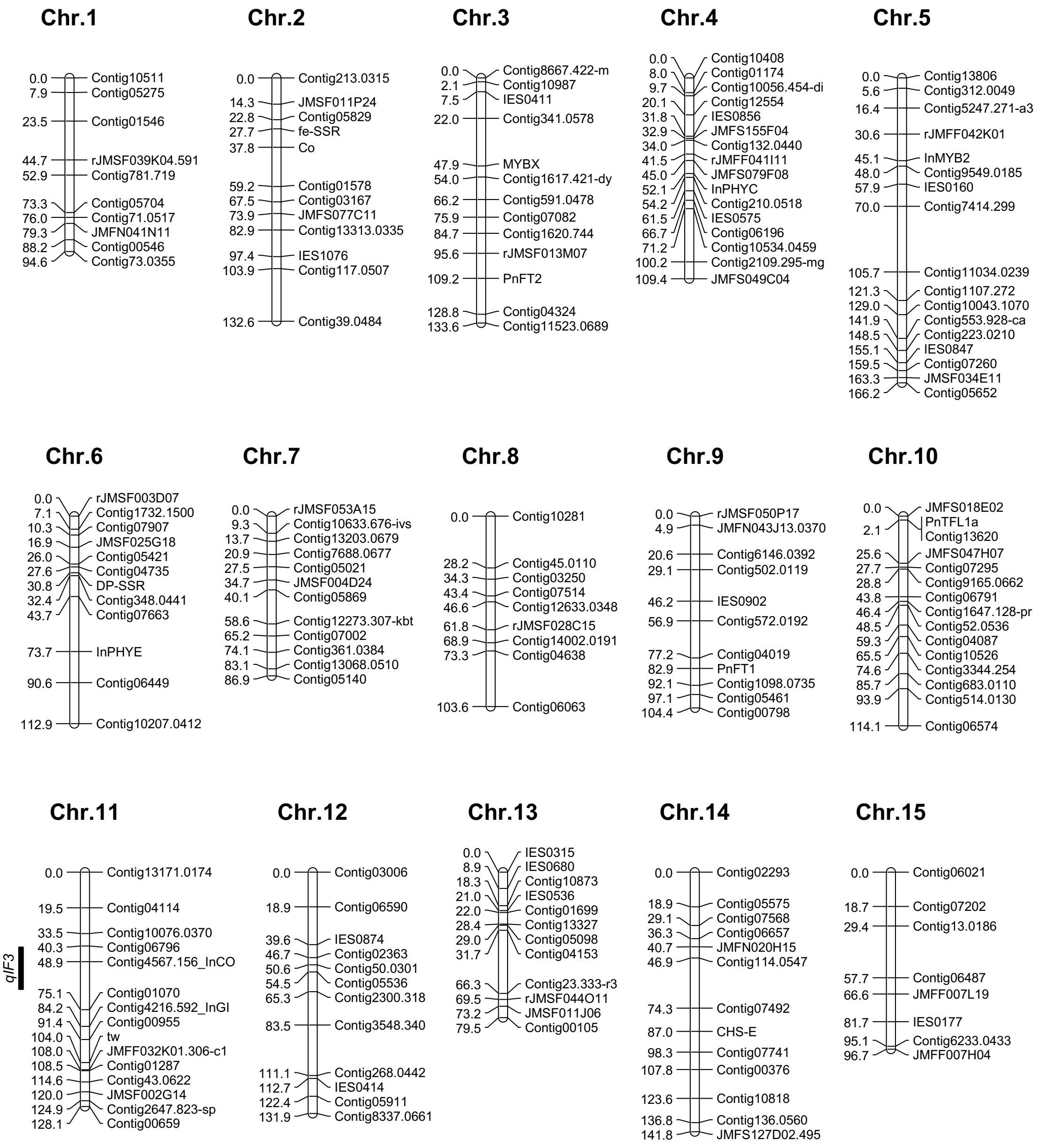


**Supplemental Fig. 2.** Genetic linkage map of $F_2$ population derived from the cross between Q65 × TKS and cultivated in 2012. The marker distance (cM) is presented on the left side of each chromosome. The marker names are shown on the right side. A vertical black bar shows a QTL region for flowering days with 99% creditable intervals.
