## Supplemental Fig. 3 for "Frame-shift mutation of *InCO* might cause early flowering of Japanese morning glory and might have contributed to northward expansion"

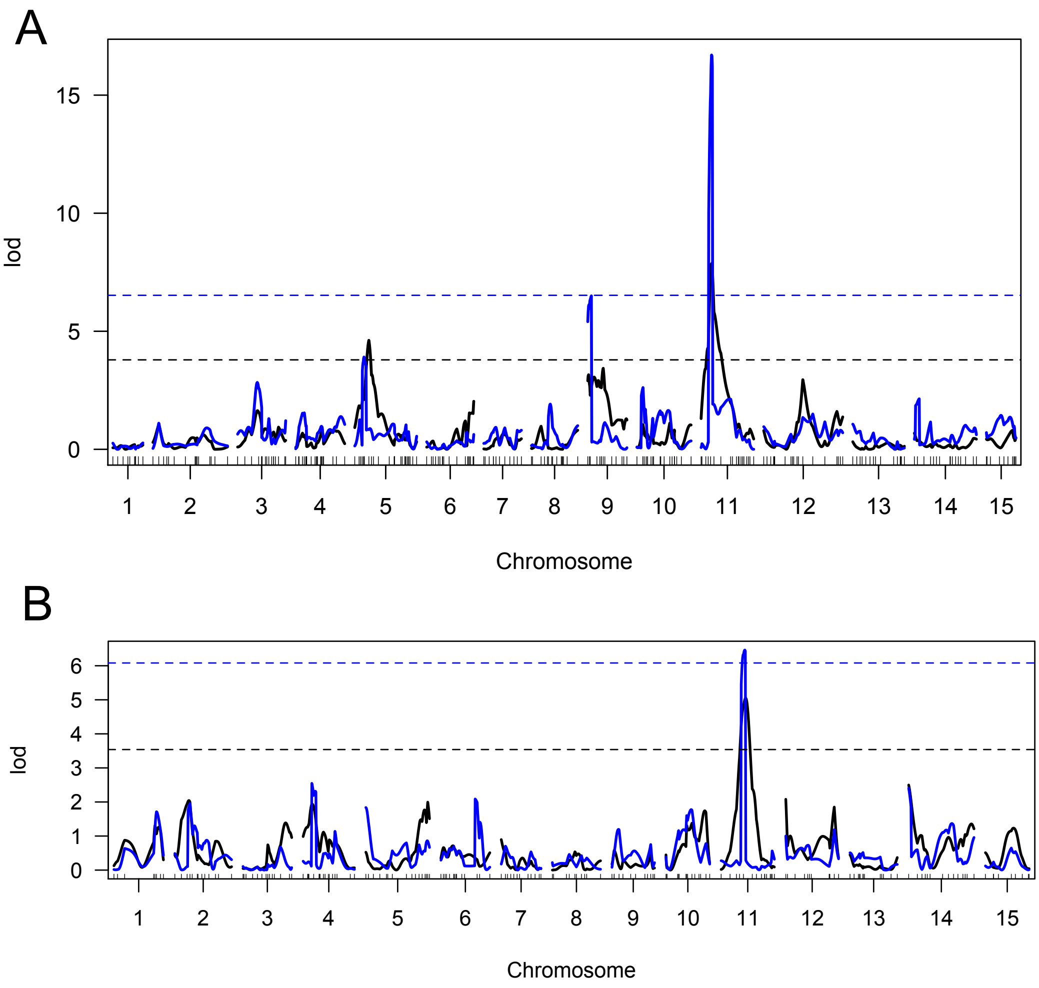


**Supplemental Fig. 3.** The LOD curves of QTL analyses for days from sowing to flowering in $F_2$ populations derived from the cross, Q65 × TKS.

A and B show LOD curves of the $F_2$ populations sown respectively on May 20, 2011 and on June 7, 2012. Black lines and blue lines respectively show the results of interval mapping (IM) and composite interval mapping (CIM). Black and blue horizontal dashed lines respectively show the permutation-test significance threshold ($n$ = 1000, $P$ = 0.05) of IM and CIM.
