## Supplemental Fig. 4 for "Frame-shift mutation of *InCO* might cause early flowering of Japanese morning glory and might have contributed to northward expansion"

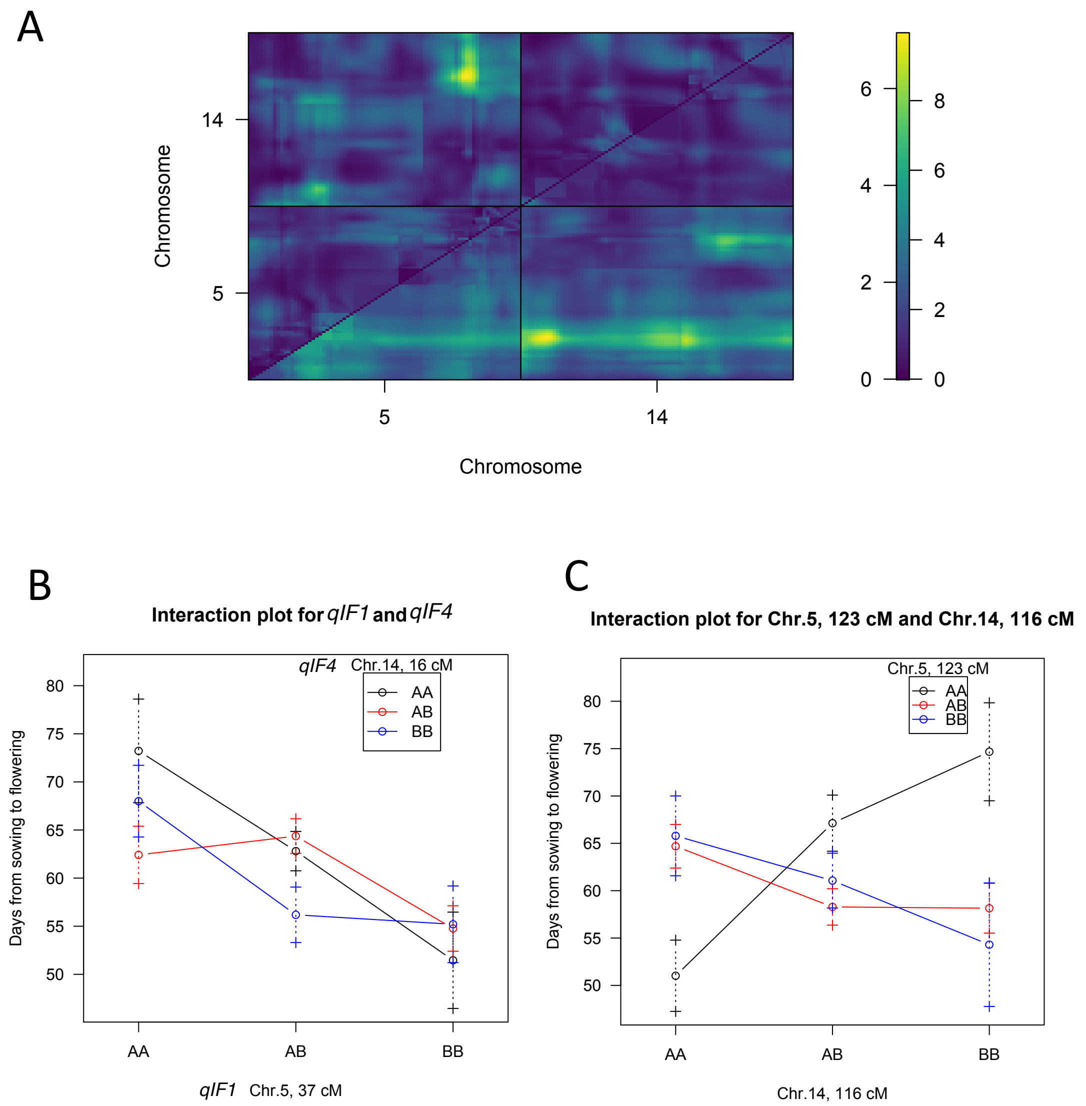


**Supplemental Fig. 4.** Interaction between QTLs located on chr. 5 and chr.14. (A) A plot of a two-dimensional, two-QTL genome scan between chr. 5 and chr.14. The upper-left triangle contains the epistasis LOD scores ("int"). The lower-right triangle contains the LOD scores for the full model ("full") in R/qtl. The significance threshold ($P = 0.05$) of LOD scores calculated through a permutation test ($n = 1000$) for the full model and for the epistasis are, respectively, 9.44 and 6.39. (B, C) Plots of days from sowing to flowering against genotypes at two pairs of loci on chr. 5 and chr. 14. (B) Plots of two loci at 37 cM on chr. 5 (*qIF1*) and at 16 cM on chr. 14 (*qIF4*). (C) Plots of two loci at 123 cM on chr. 5 and 116 cM on chr. 14.
