## Supplemental Fig. 5 for "Frame-shift mutation of *InCO* might cause early flowering of Japanese morning glory and might have contributed to northward expansion"

```
TKS     ACGGTTGTAGCAGTGCAGAACGAAAGAAAGAAGACTAAGTGCGTGGCAATGCACATCCAA 60
Q65     ACGGTTGTAGCAGTGCAGAACGAAAGAAAGAAGACTAAGTGCGTGGCAATGCACATCCAA 60
        ************************************************************

TKS     ACACGTATACTTTCTGCCTTGGATAAACATGCTTTACCAATGAAAATTATTAGCAATAAG 120
Q65     ACACGTATACTTTCTGCCTTGGATAAACATGCTTTACCAATGAAAATTATTAGTAATAAG 120
        ***************************************************** ******

TKS     CATTTCTACATCAGATACATTTTTTGACATTGAAAGCTCACACAAACCTTTTGGAGGCCA 180
Q65     CATTTCTACATCAGATACATTTTTTGACATTGAAAGCTCACACAAACCTTTTGGAGGCCA 180
        ************************************************************

TKS     TTTACATAATTAAAAAGCTTTTGAGCTATAGAAAAAGACGTTTCAATCTGCATTTAAGGG 240
Q65     TTTACATAATTAAAAAGCTTTTGAGCTATAGAAAAAGACATTTCAATATGCGTTTAAGGG 240
        *************************************** ******* *** ********

TKS     GTTTAAATTGATTTTTTTTTTTTTTGATAAATTTTT------------------------ 276
Q65     GTTTAAATTGATTTTTTTTTTTTT-GATAAACTTTTCATTCCTTTTTTTTTTTTGAAAAC 299
        ************************ ****** ****

TKS     ------------------------------------------------------------ 276
Q65     ATTTTTTTATATTCATTTTTTTTTTTTGAATGCTACTAGCTCTATTACAATGTAGTATCT 359

TKS     ------------------------------------------------------------ 276
Q65     GTTCGTATACTTTCTCAACCTACTGAAACACAAAGAGCTAATAACGCCTCCATTGAGGCT 419

TKS     ------------------------------------------------------------ 276
Q65     CGAACTCACAACCTTTTAAATGGAAATCCAACCCGGGTACCATTGGACCACATAGTCCTG 479

TKS     -----------TGTTCCTTATTCTTACTCCTATATACTTTTACATATATTTATTTAAGAA 325
Q65     GTATTTTTTTATATTCCTTATTCTTACTCTTATATACTTTTACACATATTTATTTAAGAA 539
                   * **************** ************** ***************

TKS     AAAAGTTCCAATAGACACCTCAACTCATCAATTTGATGTAATTGAGTTGATTAGCCAATT 437
Q65     AAAAGGTCAAATAAA-ACATCAAATTATTA-TTTGATGCAATTGAGTTGATTAGCCAATT 597
        ***** ** **** * ** **** * ** * ******* *********************

TKS     AACTTTAGATAATCAATACAAAGTGACATTTTAAATTTTTAATTCAAATAAATTAATTTA 497
Q65     AACTTTAAATAATTAGTATAAAGTGACATTTTAA--------TTCAAATAAAGTAATTTA 649
        ******* ***** * ** ***************        ********** *******

TKS     AAAATTTAAAAATGTTTCTTAGAGATGCTAGCCACAGCTTCCCCGTTGGACGAAGAAGTC 557
Q65     AAAATTAAAAAATATTTCTTAGAGAAGCTAGTCATGGCTTCCCCGTTGGACGAAGAAGCC 709
        ****** ****** *********** ***** **  ********************** *

TKS     CTTCTTCATTCAAATCTGGATAGAGAAGCAGACAATAATATATTTTCTTTTCAAAAAAAA 617
Q65     CTTCTTCATCCAAATCTGAATAGAGAAGTAGACAAAAATATATTTTCTT---AAAAAAAA 766
        ********* ******** ********* ****** *************   ********

TKS     AATCATTTGAAAAAGCTTACACCGTCAATGCATTTTATTGATTTCTACCTTGAGTTTACC 677
Q65     AATCATTTGAAAAAACTTACCACGT-----CATTTTATTGATTTCTACCTTTAGTTTATC 821
        ************** *****  ***     ********************* ****** *

TKS     AGCCAATTGACTTAATAACCCTTAATTACATCAATTTGATCAACTATTAGTTTAATTGTA 737
Q65     GGCCATTTGATTTAATTACA-TCAATTTGAGTCATTTGACCAACTATTACCTTAATTGTA 880
         **** **** ***** **  * ****  *   ****** *********  *********

TKS     TGAATTTGACGAGTTAAAGTGTTCATTTTTATAATTTAAGAAGCTAAGTTATTTGGCCTA 797
Q65     TCAATTTGACGAGTTAAAGTGTTCATTTTTATAATTTAAGATGCTAAGTTATTTTGCCTA 940
        * *************************************** ************ *****

TKS     TAATTTGAGAGCTTAGCTATATGATCTTCTCCTTTTTATTTATATTTATAGTGTATATAT 857
Q65     TAATTTGAGAGCTTAGCTATTTGATCTTCTCCTTTTTATTTATATTTATGGAGTATATAT 1000
        ******************** **************************** * ********
```

```
TKS   ACCCGAATATTACTTTTAATTATAAAACTAGTAAACAGTTTATATTTATTCAGAAAAGTT 917
Q65   ACCTGAATATTACTTTTAATTATAAAACTAGTAAACAATTTATATTTATTCAGAAAAATA 1060
      *** ********************************* ******************* *

TKS   ACAAACATACCCTATATTTTTTATATTTTATGTTTT--TTTTTAT-TATTTTTATTTCCA 974
Q65   ACAAACGTACTCTATATTTGTTATATTTTGTGTTTTGATTCCCATATATATATATATATA 1120
      ****** *** ******** ********* ******  **   ** *** * *** *  *

TKS   ------------------------------------------------------------ 974
Q65   TATATATATATATATATATATATATATATATATATATATATATATATATATANNNNNNNNNN 1180

TKS   -------------------------------------AATATATATATATATATATATAT 997
Q65   TATATATATATATATATATATATATATATATATATATAATATATATATATATATATATAT 1240
                                           ***********************

TKS   ATATATATATATATATATATATATATATATATATATATATATATATATATAT-------- 1049
Q65   ATATATATATATATATATATATATATATATATATATATATATATATATATATATATATAT 1300
      ****************************************************

TKS   -----------------------GTTATCTATTTTC--------CAGATCGCATCAAGGA 1078
Q65   ATATATATATATATATAATTATTGTTATCTATTTCCATGCTACCTAGACCACATCAAGGA 1360
                             *********** *         *** * *********

TKS   AAGCTCATAAGAACAGGTTCATACCAGCGCTCACACGTCGTCTATTAATTATGAAAAAGG 1138
Q65   AAACTCATAAGAACAGGTTCATACTAGGACTCACACTTCGTCTATTAATTAGAAAAAGTG 1420
      ** ********************* **  ******* **************  ****  *

TKS   CCAAATAAATCATTAAATTTTTACACTTTGTACAATAGAACCATCAAATAAAAAAGTG-- 1196
Q65   TAAAATAGACCACTGAACTTTTACAATTTTTATAATTGAATTATAAAATTAAAAAGTGCA 1480
        ***** * ** * ** ******* *** ** *** ***  ** **** ********

TKS   ---AGGCCATTAAAAAAATAAAAATTGTGCAAATAACATTCTTTACAAATTATTTCTAAG 1253
Q65   ATTGGAGCATCAAAAAAATAAAAATTGTGCTAATAACATTATTTACAAATTACTTCTATG 1540
          *  *** ******************* ********* *********** ***** *

TKS   TTTCTAGTAAAATTTGATGTCATAATGTTAAATTAAATATAAAAATAATT---------- 1303
Q65   TTTCTGGTAAAAGTTGATGTTATAATGTTAAATTATATATAAAAATTATTTTAATTATTA 1600
       ***** ****** ******* ************** ********** ***

TKS   --------------------AAAAAAAAATCAAAAGAAAGTTCATCTACTGTGGATAAAG 1343
Q65   TTTAAAAATATTTTTTAAAAAAAATAAATTCAGAAAAAATTTCATCTGTTGTGAATGAAA 1660
                          **** *** *** ** *** *******  **** ** **

TKS   GGAAAGGCCGTCATGGCTGCTCCCACCTTCGTC---------------------CAA--- 1379
Q65   GGAAAAGTAGCTATGACTGCCCCTACCTTCGTCTGCTATGGATGGAGGGGGAGGCAACCA 1720
      ***** *  *  *** **** ** *********                     ***

TKS   -------------TTTGAACGAAGGGGAGGCAACCATGGCTGCTCCCTCCTTCATCCAAA 1426
Q65   TGGTTGCCCCACCTTTGGACGAAGGGGAGGCAGCCATGGCTGCTCCCTCCTTCATCCAGA 1780
                   **** ************** ************************* *

TKS   TCGGATAAAGGAGGGGGCAACCATGGCTGCCTCCATCCTCGTCCAAAACAAAGGTGAGGG 1486
Q65   TCGGACGAAGAAGGGGGCAACCATAGCTGCCTCCATCCTCGTCTAGGACAAAGGTGGGGA 1840
      *****  *** ************* ****************** *  ********* **

TKS   CAACCATGGTTATCATTCCCTTCATCCACAACAAACGAAGGTGTAGGCAATCATGCCCTT 1546
Q65   CAGTAATGGTTGTC--TCCCT------------GATGAAAGGGTAGGTAGTCGTGCCCTT 1886
      **   ****** **  *****             * *** * ***** * ** *******
```

```
TKS    ACACCTT---TTAAAATTAAATTTTTT-------------TTTAAAATAATAATTAAAAT 1590
Q65    CCACCTTCTTTTGAAATTAAATTTTTTGTTTTTAATTTTTTTTAAAATAATAATAATAAT 1946
        ******   ** **************             ************** * ***

TKS    AAT-----------TTATATATATACACCATA---ATAATTTAACATTATGACATTATAT 1636
Q65    AATAATAATAATAATAATTAAAATAATTTATATATATAATTTAACATTATGACATTATAT 2006
       ***           * **  * ***    ***   *************************

TKS    TTAATAGAAACTTGAAAATAACTTATAATAAAATACTATTTGTATAATTTTTGAGTTCTT 1696
Q65    TTAATAGAAACTTGAAAATAACTTATAATAAAATACTATTTGTATAATTTTTGAATTCTT 2066
       ****************************************************** *****

TKS    AATAATTTAATTACATATTTTTTTTGTTCAATGATCAATTACACACAATGTAAAAGTTCA 1756
Q65    AATAATTTAATTACATATTTTTTTTGTTCAATGATCAATTACACACAATGTAAAAGTTCA 2126
       ************************************************************

TKS    ATGGCCTATATGACATTTTTTTCACTAATTTGATGTGAAGAACGTTTCTCCCAGAGTGTG 1816
Q65    ATGGCCTATATAACATTTTTTTCACTAATTTGATGTGAAGAACGTTTCTCCCAGAGTGTG 2186
       *********** ************************************************

TKS    GGCAGCAAACGGATTCCAAAAGCCAAAGCCATTGACAGGACAGGTCTGTAAAACAGAATC 1876
Q65    GGCAGCAAACGGATTCCAAAAGCCAAAGCCATTGACAGGACAGGTCTGTAAAACAGAATC 2246
       ************************************************************

TKS    TGGGTCAGCGGGACCCCGCGGCTGGCCCCACCAGAATCACAGCCAGCCACAAGATAATAC 1936
Q65    TGGGTCGGCGGGACCCCGCGGCTGGCCCCACCAGAATCACAGCCAGCCACAAGATAATAC 2306
       ****** *****************************************************

TKS    AACCTTTTGCAACCCGCTAATATCGCACCACGTGTCAGCTCTGGATTCTCCTCCCACTCT 1996
Q65    AACCTTTTGCAACCCGCTAATATCGCAACACGTGTCAGCTCTGGATTCTCCTCCCACTCT 2366
       **************************** *******************************

TKS    CCCTTCCAACTTGCTCACTGCTACCACAAAGTCATAAAAAGCATAGCTGCAGGACTCGGT 2056
Q65    CCCTTCCAACTTGCTCACTGCTACCACAAAGTCATAAAAAGCATAGCTGCAGGACTCGAT 2426
       ********************************************************** *

TKS    CACTCTAAATAAATACGTTTGAGTGTGTGTAGTCTTACGTGTGTGCGAGGAAACACTCAA 2116
Q65    CACTCTAAATAAATACGTTTGAGTGTGTGTAGTCTTACGTG----GGAGGAAACACT-AA 2481
       *****************************************     *********** **
```

**Supplemental Fig. 5.** DNA sequence alignment of the 5' flanking region of *InCO* between TKS and Q65. Yellow highlights denote a SINE-like sequence. Light blue highlights denote target site duplications. A bent arrow indicates a putative transcript-start site. TKS, Tokyo Kokei standard.
