## Supplemental Fig. 6 for "Frame-shift mutation of *InCO* might cause early flowering of Japanese morning glory and might have contributed to northward expansion"

```
TKS    MLKEESCEVLDLDVTIGSSSGSRSGNKQNWARVCDICRSAACSVYCRADLAYLCGGCDAR   60
Q65    MLKEESCEVLDLDVTIGSSSGSRSGNKQNWARVCDICRSAACSVYCRADLAYLCGGCDAR   60
       ************************************************************

TKS    VHGANTVAGRHERVLVCEACESAPATVICKADAASLCAACDSDIHSANPLARRHHRVPIL   120
Q65    VHGANTVAGRHERVLVCEACESAPATVICKADAASLCAACDSDIHSANPLARRHHRVPIL   120
       ************************************************************

TKS    PISGTLYGPPTSNPCRESSMMVGLTGDAAEEDNGFLTQDAEETTMDEDEDEAASWLLLNP   180
Q65    PISGTLYGPPTSNPCRESSMMVGLTGDAAEEDNGFLTQDAEETTMDEDEDEAASWLLLNP   180
       ************************************************************

TKS    NPNPNPNP----VKSNNSTNMCKGGNNNNN---EMSCAVEAVDAYLDLAEFSSCHNNLFE   233
Q65    NPNPNPNPNPNPVKSNNSTNMCKGANNNNNNNNEMSCAVEAVDAYLDLAEFSSCHDNLFE   240
       ********    ************.*****   **********************:****

TKS    DKYSINQQQNYSVPQRNMSYRGDSIVPNHGKNQFHYTQGLQQHNHHAIFNCKEWNMRILT   293
Q65    DKYSINQQQNYSVPQRNMSYRGDSIVPNHGKNQFHYTQGLQQHNHHRNFQLQGMEYENFN   300
       **********************************************  *: :  : . :.

TKS    RDMVIQHPSVTQSPFLPWMLVLFQSLP---------------------------------   320
Q65    TGYGYPASISHTVSISSMDVGVVPESTLSDASISHSRASKGTIDLFSGPPIQMPPQLQLS   360


TKS    ------------------------------------------------------------
Q65    QMDREARVLRYREKKKTRKFEKTIRYASRKAYAETRPRIKGRFAKRTDVDTEVDQIFYAP   420
                                CCT domain

TKS    -------------
Q65    LMAESGYGIVPSF   433
```

**Supplemental Fig. 6.** An amino acid sequence alignment of the InCO/IhCO protein, encoded by the transcript variant *InCO* (si) of the TKS allele and the Q65 allele. Sequences were aligned with Clustal W (Ver. 1.83, 2003).
